# Stomatal Complex Ionomes Explain Divergent Gas Exchange Responses to Salinity in Maize and Faba Bean

**DOI:** 10.64898/2026.08.10.743902

**Authors:** Xudong Zhang, Guanghui Wei, Christian Zörb

## Abstract

Salinity tolerance is commonly associated with whole leaf Na⁺ exclusion and maintenance of K⁺ homeostasis, but whether spatial ion partitioning among functional leaf compartments contributes to stress adaptation remains unclear. Here, we investigated the relationship between bulk leaf and stomatal complex ionomes and gas exchange performance under salinity using two contrasting genotypes in both maize and faba bean crops. Maize generally maintained higher photosynthesis and stomatal conductance than faba bean under salt stress, which was associated with lower Na⁺ accumulation, stronger K⁺ retention and distinct ion partitioning patterns between bulk leaf tissue and the stomatal complex. Enrichment analysis revealed that stomatal complex ion composition provided information beyond bulk leaf ion concentrations, with Na⁺ and Cl⁻ showing distinct distribution patterns associated with photosynthetic performance. Integrating physiological and ionomic traits further demonstrated that stomatal-complex ion traits captured additional variation in salinity responses. These findings identify the stomatal complex as a functionally distinct ionomic compartment and reveal compartment-specific ion partitioning as an important mechanism underlying species-specific salinity tolerance.

## 1 Introduction

Salinity is one of the major abiotic stresses limiting plant productivity worldwide (Zörb et al., 2019). Increasing soil salinization caused by climate change, irrigation practices and land degradation poses a serious challenge to sustainable crop production (Singh, 2022; Tarolli et al., 2024). Excessive accumulation of Na⁺ and Cl⁻ disrupts cellular ion homeostasis, reduces K⁺ availability, impairs photosynthesis and ultimately limits plant growth and yield (Munns and Tester, 2008; Van Zelm et al., 2020). Plants have evolved diverse mechanisms to tolerate salinity, including Na⁺ exclusion, vacuolar sequestration, maintenance of K⁺ homeostasis and regulation of ion transport across tissues (Deinlein et al., 2014). Understanding how these mechanisms operate at different organizational levels remains essential for improving crop salt tolerance.

Most studies on salinity tolerance have focused on bulk tissue ion concentrations, particularly leaf Na⁺, Cl⁻ and K⁺ concentrations (Flowers et al., 2015). Reduced Na⁺ accumulation and maintenance of K⁺ levels are generally considered key indicators of salt tolerance (Almeida et al., 2017; Wu, 2018). However, bulk leaf ion measurements represent the integrated outcome of ion uptake, transport, storage and compartmentation across different cell types. Therefore, similar ion concentrations at the tissue level may have different physiological consequences depending on their spatial distribution within the leaf. Recent advances in ion imaging and cell type-specific analyses have revealed substantial heterogeneity in elemental distribution, emphasizing the importance of understanding ion partitioning among functional compartments (Conn and Gilliham, 2010; Fernández-Ramírez et al., 2026).

Stomata represent a critical interface between ion regulation and plant performance because they directly control gas exchange, water loss and carbon assimilation (Lawson and Vialet-Chabrand, 2019). Stomatal function depends on dynamic regulation of osmotic and ionic conditions within guard cells and surrounding epidermal tissues (Hedrich and Shabala, 2018). The ions K⁺ and Cl⁻ contribute to stomatal movement, while ion imbalance caused by salinity can affect stomatal regulation and photosynthetic activity (Kollist et al., 2014; Lawson et al., 2014). However, how salinity alters ion composition within the stomatal complex and whether this local ionome contributes to species-specific salt tolerance remain largely unknown.

This knowledge gap is particularly relevant when comparing plant species with contrasting salt responses. Maize, a C₄ monocot cereal, generally exhibits higher salt tolerance than many C₃ dicot legumes, including faba bean, reflecting differences in photosynthetic performance and stomatal complex morphology (Page et al., 2021). However, the mechanisms underlying these differences are not fully understood. Comparative analysis of species with contrasting physiological strategies provides an opportunity to determine whether salinity tolerance depends mainly on overall ion exclusion or on the spatial regulation of ions within specific functional compartments.

Here, we investigated how bulk leaf and stomatal complex ionomes contribute to divergent gas exchange responses under salinity using contrasting genotypes in maize and faba bean. We hypothesized that (*i*) salinity tolerance is associated with species-specific ion partitioning between the bulk leaf and stomatal complex, (*ii*) stomatal-complex ion traits provide additional information beyond bulk leaf ion composition, and (*iii*) Na⁺ and Cl⁻ exhibit distinct partitioning patterns linked to gas exchange regulation. To test these hypotheses, we integrated gas exchange measurements, bulk leaf ion profiling, stomatal complex ion analysis and ion enrichment indices. Our study identifies the stomatal complex ionome as an additional functional layer of ion regulation and provides new insight into the mechanisms underlying species-specific salinity adaptation.

## 2 Material and Methods

### 2.1 Plant cultivation

Two contrasting maize (*Zea mays* L.) genotypes, LG30222 (LG c/o Limagrain GmbH) and ES-Metronom (Euralis Saaten GmbH) (Zhang et al., 2019), and two contrasting faba bean (*Vicia faba* L.) genotypes, Fuego and Scoop (Norddeutsche Pflanzenzucht Hans-Georg Lembke KG, Hohenlieth, Germany) (Franzisky et al., 2019), were cultivated hydroponically in climate-controlled growth chambers (WEISS HGC1014, Heuchelheim, Germany). Plants were grown under a 14 h light and 10 h dark photoperiod at 22/18 °C day and night. Relative humidity was maintained at 80 to 90% for maize and approximately 80 to 60% for faba bean, with light intensities of approximately 500 and 300 μmol photons m⁻² s⁻¹ at canopy level, respectively. Cultivation followed previously described protocols with minor modifications (Franzisky et al., 2025; Zhang et al., 2020).

Seeds were soaked in aerated 0.5 mM CaSO₄ solution for 24 h before germination in moist quartz sand. Approximately 12 days after germination, seedlings were transferred to one-quarter-strength aerated nutrient solution, which was gradually increased to full strength over four days. The maize nutrient solution contained 0.2 mM KH₂PO₄, 1.0 mM K₂SO₄, 2.0 mM Ca(NO₃)₂, 0.6 mM MgSO₄, 0.2 mM Fe-EDTA, 10 μM NaCl, 10 μM H₃BO₃, 2.0 μM MnSO₄, 0.5 μM ZnSO₄, 0.2 μM CuSO₄, 0.1 μM CoCl₂ and 0.05 μM (NH₄)₆Mo₇O₂₄ (Zhang et al., 2020). The faba bean nutrient solution contained 0.1 mM KH₂PO₄, 1.0 mM K₂SO₄, 2.0 mM Ca(NO₃)₂, 0.5 mM MgSO₄, 0.126 mM Fe-EDTA, 10 μM NaCl, 10 μM H₃BO₃, 2.0 μM MnSO₄, 0.5 μM ZnSO₄, 0.2 μM CuSO₄, 0.1 μM CoCl₂ and 0.05 μM (NH₄)₆Mo₇O₂₄ (Franzisky et al., 2025). Nutrient solutions were continuously aerated and renewed twice weekly.

After acclimation to full-strength nutrient solution, maize and faba bean plants were subjected to the same five treatments consisting of a control (1 mM NaCl), 50 mM NaCl, 100 mM NaCl, 50 mM Na₂SO₄ and 50 mM CaCl₂. To minimise osmotic shock, salt concentrations were increased gradually over three days by adding one-third of the final concentration each day until the target concentration was reached.

Gas exchange measurements were performed on the fifth fully expanded leaf between 4 and 6 h after the onset of the light period for seven consecutive days following full salt exposure. Plants were harvested on the eighth day after full salt treatment. Leaf epidermal strips from the fifth leaf were mounted on copper grids and cryo-preserved in liquid nitrogen for cryogenic scanning electron microscopy coupled with energy-dispersive X-ray spectroscopy (cryo-SEM-EDX). The remaining leaf tissue was frozen, ground and freeze-dried for ionomic analysis by inductively coupled plasma optical emission spectroscopy (ICP-OES). Each treatment consisted of three pots with three plants per pot.

### 2.2 Gas exchange

Gas exchange parameters, including net photosynthetic rate (A), transpiration rate (E), and stomatal conductance (gs, relative to CO₂), were measured using a portable photosynthesis system (GFS-3000, Heinz Walz GmbH, Effeltrich, Germany). The measuring cuvette (2.5 cm² cross-sectional area) was maintained at 400 ppm CO₂, 15,000 ppm H₂O, 25 °C air temperature, 1000 μmol m⁻² s⁻¹ photosynthetically active radiation (PAR), and a gas flow rate of 750 μmol s⁻¹. At least five measurement spots were recorded from five independent plants per treatment.

### 2.3 Ionome analysis

For ionome analysis, approximately 50 mg of freeze-dried leaf material was subjected to microwave-assisted pressure digestion with concentrated nitric acid (65% HNO₃) in quartz vessels using an UltraClave V microwave system (MLS, Leutkirch, Germany) according to established protocols (Wollmann et al., 2018). The digested samples were analysed by ICP-OES (Agilent 5110, Agilent Technologies, Santa Clara, CA, USA) for Na, K, Ca, Mg, P and S. Chloride (Cl) was quantified separately by ion chromatography (IC) with suppressed conductivity detection. Quantification was performed using external calibration with certified multi-element and single-element standards. Samples with Na concentrations below the detection limit (<400 ppm) were assigned a value of half of the detection limit (200 ppm) for statistical analysis and visualization.

### 2.4 Microscopic observation and semi-quantitative surface elemental analysis

Cryo-SEM-EDX was used to analyse the surface-resolved elemental composition of stomatal complexes. Epidermal strips from the fifth fully expanded leaves used for gas exchange measurements were mounted on copper grids and plunge-frozen in liquid nitrogen to preserve native elemental distributions. Samples were transferred into a cryo-preparation chamber (Quorum Technologies) connected to a Zeiss Crossbeam 550 SEM without freeze fracture or sublimation treatment. Samples were sputter-coated with platinum (5 mA, 45 s; approximately 35 nm) and analysed using an Oxford Instruments X-MaxN 150 SDD detector controlled by AZtec software (v5.1) (Franzisky et al., 2025).

Measurements were performed at 8 kV accelerating voltage, 1 nA beam current, 2126× magnification and 5.5 mm working distance. Regions of interest corresponding to the entire stomatal complex were manually selected based on SEM morphology. In faba bean, the stomatal complex included the stomatal pore and the surrounding pair of guard cells, whereas in maize it additionally included the surrounding pair of subsidiary cells. The stomatal complex was treated as a single analytical unit without separating individual component cells. At least 10 stomatal complexes per treatment were analysed from five epidermal peel samples, with each spectrum acquired for ≥5 min. Elemental abundances were expressed as atomic percentages (atom%) and normalized to the sum of detected biological elements for semi-quantitative comparison.

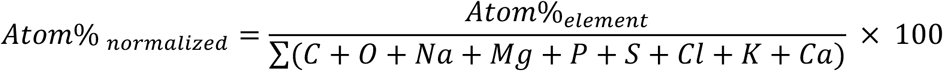

Detailed information regarding EDX signal interpretation, surface detection depth, element selection criteria, spectral validation and below-detection-limit handling is provided in Supplementary Method S1. Representative EDX spectra across all genotype–treatment combinations are shown in Figure S1.

### 2.5 Relative ion enrichment patterns in stomatal complex surface regions

Leaf ion concentrations measured by ICP-OES (mg g⁻¹ DW) and stomatal complex (StC) elemental signals obtained by cryo-SEM-EDX (atom%) were analysed separately to describe bulk leaf ion accumulation and stomatal complex-associated ion distribution, respectively. To quantify relative ion partitioning between the stomatal complex and whole leaf, an enrichment index was calculated as the log₁₀ ratio between normalized StC and leaf ion responses:

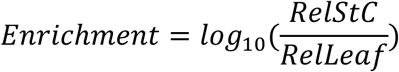

where relative ion responses were calculated by normalizing each treatment value to the corresponding genotype-specific control. Because some ion measurements contained zero or near-zero values, a small pseudocount (ε) was added before ratio calculation and log₁₀ transformation (Lambert et al., 1991; Martín-Fernández et al., 2003). The ε value was defined as half of the minimum non-zero value across all samples. Sensitivity analysis using alternative ε values ranging from 0.5× to 2× of this value confirmed consistent enrichment patterns across genotypes and treatments (Figure S2).

The enrichment index is a unitless measure that enables direct comparison between EDX-derived stomatal complex surface elemental signals and ICP-OES-derived bulk leaf ion concentrations. Positive values indicate relative ion enrichment in the stomatal complex compared with the whole leaf, whereas negative values indicate relative depletion. Thus, the enrichment index reflects the relative partitioning of ions between stomatal complex surface regions and bulk leaf tissues.

For statistical analysis, enrichment indices were calculated for each stomatal complex measurement using the corresponding mean leaf ion concentration from the same genotype and treatment. Mean enrichment values ± standard error were used to evaluate treatment- and genotype-dependent differences in relative ion partitioning.

### 2.6 Correlation between ion enrichment patterns and net photosynthetic rate

All treatments were included in the correlation analysis to capture the overall relationship between ion partitioning and photosynthetic responses. Although some treatments showed minimal variation in specific ions (for example, Na in CaCl₂ treatment and Cl in Na₂SO₄ treatment), these data points were retained because they represent baseline ion conditions relative to the control and contribute to the integrated assessment of physiological responses. Correlations were analysed between net photosynthetic rate (A) and Na and Cl enrichment indices. Linear regressions were fitted separately for each genotype and species, and regression slopes and Pearson correlation coefficients (r) were used to evaluate differences in ion–photosynthesis relationships among genotypes and species. Because stomatal complex elemental signals obtained by cryo-SEM-EDX were derived from pooled epidermal samples rather than individual plants, enrichment indices could not be directly matched with individual gas exchange measurements. Therefore, correlations were performed using treatment means, representing the integrated ionic and physiological responses of each genotype under different salt conditions.

### 2.7 Principal component analysis

To characterize genotype-specific physiological and ionomic responses under salt stress, principal component analysis (PCA) was performed using treatment-averaged values of gas exchange parameters, leaf and stomatal complex elemental profiles, and Na and Cl enrichment indices. PCA was used to visualize multivariate relationships among traits and the separation of genotypes based on their integrated physiological and ionic responses.

### 2.7 Statistical analysis

Data were analysed using two-way analysis of variance (ANOVA) to assess the effects of genotype, treatment and their interaction (genotype × treatment), followed by Fisher’s least significant difference (LSD) test for multiple comparisons implemented in the agricolae package in R. Gas exchange measurements were based on five biological replicates per treatment. Leaf ionomic analyses were performed using three to four biological replicates per treatment. Stomatal complex elemental composition was determined by SEM–EDX analysis of randomly selected stomatal complexes from epidermal samples collected across five independent plants, with at least 10 stomatal complexes analysed per treatment. Ion enrichment indices were calculated as log₁₀ ratios between normalized stomatal complex elemental signals and corresponding whole leaf ion concentrations, with observations based on individual analysed stomatal complexes. Figures were generated using GraphPad Prism (version 10.2.0). Principal component analysis was performed in R using prcomp, and PCA visualizations were generated using ggplot2.

## 3 Results

### 3.1 Genotypic and species-specific gas exchange responses to salinity

To characterize physiological responses to salinity, net photosynthetic rate (A), transpiration rate (E) and stomatal conductance (gs) were measured in two maize genotypes (LG30222 and ES-Metronom) and two faba bean genotypes (Fuego and Scoop) under control and four salt treatments (Fig. 1). Two-way ANOVA revealed significant effects of genotype, treatment and their interaction for all three gas exchange parameters (P < 0.001).

**Figure 1.**
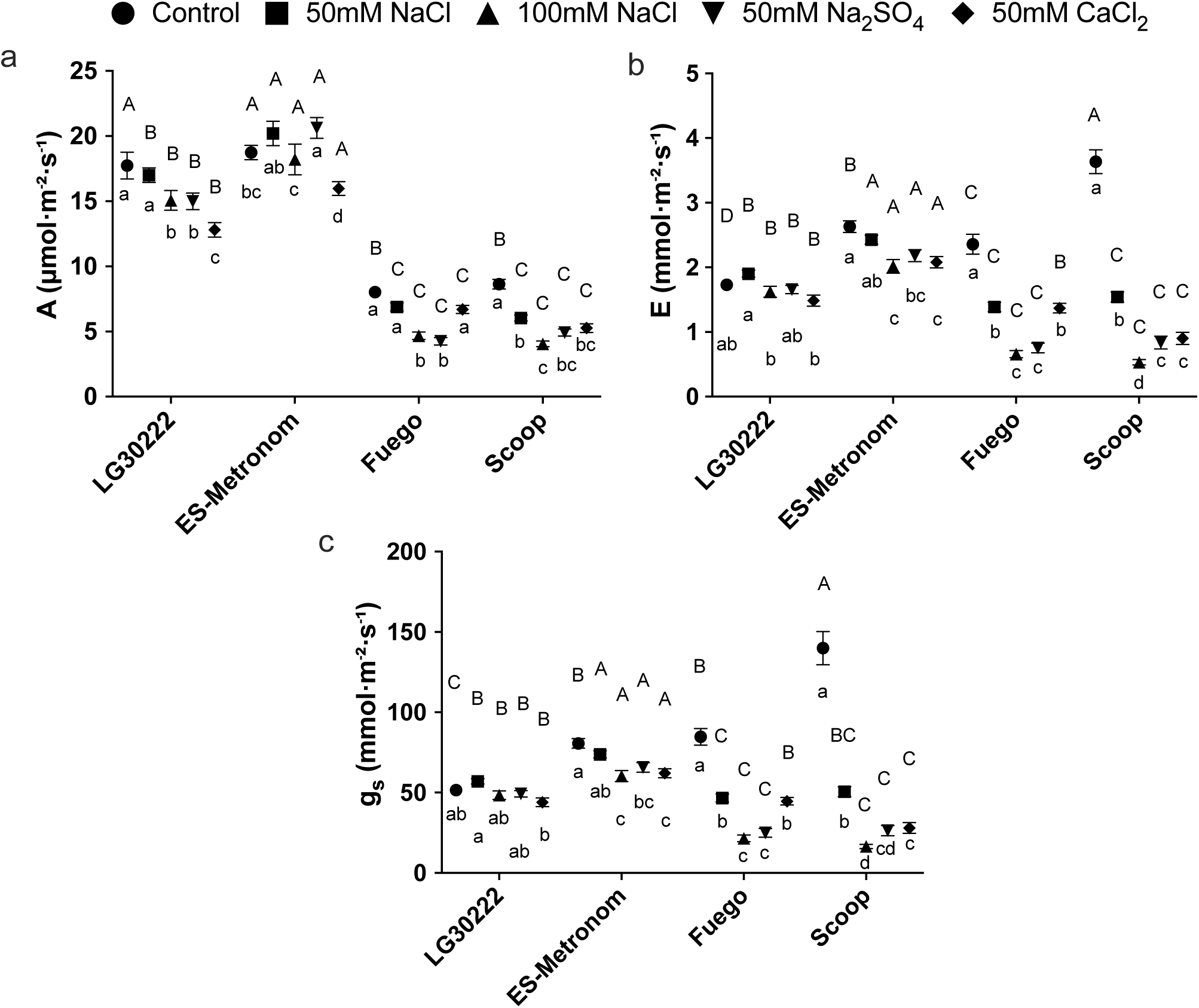
Gas exchange responses in maize and faba bean under salt stress. **a–c**, Net photosynthetic rate (A) (a), transpiration rate (E) (b), and stomatal conductance (gs) (c) in leaves of two maize genotypes (LG30222 and ES-Metronom) and two faba bean genotypes (Fuego and Scoop). Data are presented as means ± s.e. (n = 5 biological replicates per treatment). Two-way ANOVA revealed significant effects of treatment, genotype and their interaction (all P < 0.001). Different uppercase letters indicate significant differences among genotypes within the same treatment, whereas lowercase letters indicate significant differences among treatments within the same genotype (Fisher’s LSD test, P < 0.05).

Gas exchange responses showed clear differences between species across treatments. Maize maintained higher A, E and gs than faba bean under both control and saline conditions. Within maize, ES-Metronom generally showed higher gas exchange than LG30222 under salt treatments. Under 100 mM NaCl, ES-Metronom maintained higher A and gs than LG30222, with values of 18.2 versus 15.1 µmol m⁻² s⁻¹ and 60.3 versus 48.4 mmol m⁻² s⁻¹, respectively (Fig. 1a, c). Across salt treatments, maize genotypes showed relatively stable gas exchange responses, with treatment differences mainly observed in gs. Transpiration followed a similar trend, with ES-Metronom maintaining consistently higher E than LG30222 under saline conditions, whereas both maize genotypes showed only moderate reductions compared with the control (Fig. 1b).

In contrast, both faba bean genotypes showed lower gas exchange under saline conditions compared with maize (Fig. 1). Under salt treatments, A remained below 7 µmol m⁻² s⁻¹ in both Fuego and Scoop, with similar responses between the two genotypes (Fig. 1a). Transpiration decreased markedly in both genotypes under salinity, with the strongest reductions occurring under 100 mM NaCl, while CaCl₂ and 50 mM NaCl partially alleviated the decline (Fig. 1b). Stomatal responses varied among salt treatments and genotypes. In Fuego, 50 mM CaCl₂ maintained higher gs than 100 mM NaCl and 50 mM Na₂SO₄, whereas in Scoop, CaCl₂ increased gs compared with 100 mM NaCl but showed no difference from 50 mM Na₂SO₄ (Fig. 1c).

### 3.2 Leaf ion profiles in maize and faba bean under salt stress

To characterize ion accumulation patterns associated with salt responses, we quantified leaf Na, Cl, K and K/Na ratios in the four genotypes (Fig. 2). Two-way ANOVA revealed significant effects of genotype, treatment and their interaction for all measured ion traits (P < 0.001).

**Figure 2.**
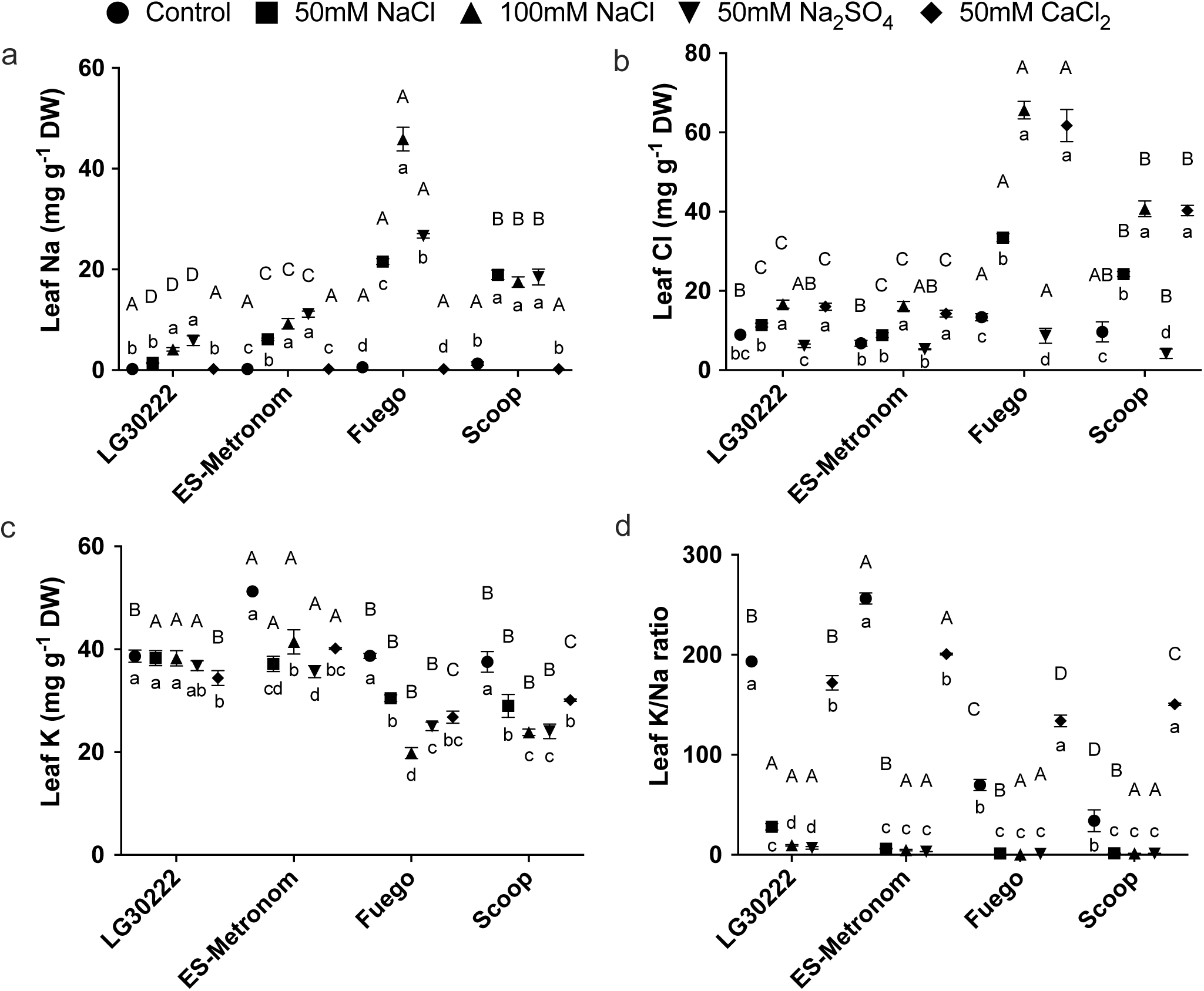
Leaf ion profiles in maize and faba bean under salt stress. **a–d**, Leaf sodium (Na) concentration (a), chloride (Cl) concentration (b), potassium (K) concentration (c) and K/Na ratio (d) in two maize genotypes (LG30222 and ES-Metronom) and two faba bean genotypes (Fuego and Scoop). Data are presented as means ± s.e. (n ≥ 3 biological replicates per treatment). Two-way ANOVA revealed significant effects of treatment, genotype and their interaction (all P < 0.001). Different uppercase letters indicate significant differences among genotypes within the same treatment, whereas lowercase letters indicate significant differences among treatments within the same genotype (Fisher’s LSD test, P < 0.05).

Leaf Na accumulation showed clear differences between species (Fig. 2a). Under NaCl and Na₂SO₄ treatments, maize accumulated substantially less Na than faba bean. Under 100 mM NaCl, leaf Na ranged from 4.1 to 9.2 mg g⁻¹ DW in maize, whereas faba bean reached 17.5 and 45.9 mg g⁻¹ DW in Scoop and Fuego, respectively. Similar differences were observed under 50 mM NaCl and 50 mM Na₂SO₄. In contrast, 50 mM CaCl₂ did not increase leaf Na accumulation, with all genotypes remaining close to control levels.

Leaf Cl accumulation displayed distinct treatment- and species-dependent patterns (Fig. 2b). Maize maintained relatively moderate Cl concentrations under NaCl and CaCl₂ treatments, whereas faba bean showed strong Cl accumulation under these conditions. The highest Cl concentrations were observed in Fuego under 100 mM NaCl and 50 mM CaCl₂, reaching 65.6 and 61.7 mg g⁻¹ DW, respectively. Under 50 mM Na₂SO₄, Cl accumulation remained low across all genotypes.

Leaf K showed contrasting responses between species (Fig. 2c). Maize maintained higher K concentrations across salt treatments, generally remaining above 35 mg g⁻¹ DW. In contrast, faba bean showed stronger K reduction under salinity, particularly under 100 mM NaCl, where K decreased to 19.8 and 23.8 mg g⁻¹ DW in Fuego and Scoop, respectively. The CaCl₂ treatment maintained higher K levels in faba bean compared with NaCl treatments at the same Cl concentration.

Consistent with changes in Na and K, K/Na ratios were strongly reduced by NaCl and Na₂SO₄ treatments but remained high under CaCl₂ (Fig. 2d). Under 100 mM NaCl, K/Na ratios ranged from 0.44 in Fuego to 9.48 in LG30222, whereas CaCl₂ maintained ratios above 130 across all genotypes.

### 3.3 Stomatal complex ion partitioning in maize and faba bean under salinity

To evaluate ion distribution within stomatal complexes, Na, Cl and K atomic percentages were quantified using stomatal complex measurements (Fig. 3). Two-way ANOVA showed significant effects of genotype, treatment and their interaction for all measured elements (P < 0.001).

**Figure 3.**
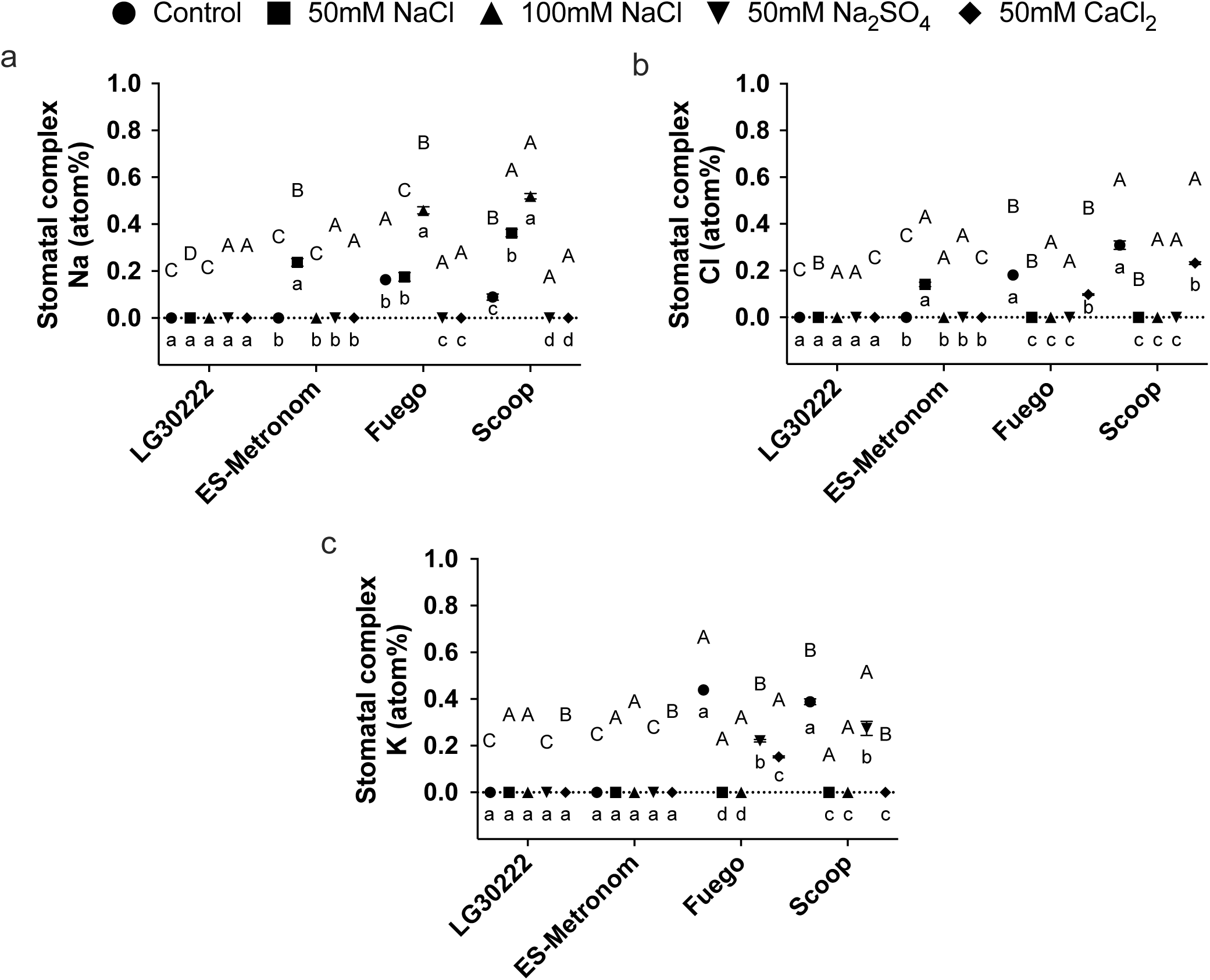
Stomatal complex ion composition in maize and faba bean under salt stress. **a–c**, Sodium (Na) (a), chloride (Cl) (b) and potassium (K) (c) concentrations in stomatal complexes, expressed as atomic percentages, in two maize genotypes (LG30222 and ES-Metronom) and two faba bean genotypes (Fuego and Scoop). Data are presented as means ± s.e. (at least 10 stomatal complexes per treatment). Two-way ANOVA revealed significant effects of treatment, genotype and their interaction (all P < 0.001). Different uppercase letters indicate significant differences among genotypes within the same treatment, whereas lowercase letters indicate significant differences among treatments within the same genotype (Fisher’s LSD test, P < 0.05).

Stomatal complex Na accumulation differed between species (Fig. 3a). Maize generally maintained low Na levels, with LG30222 remaining near zero across all treatments. ES-Metronom showed a specific increase under 50 mM NaCl, reaching 0.237 atom%. In contrast, faba bean accumulated higher Na under NaCl treatments, with maximum values of 0.519 atom% in Scoop and 0.459 atom% in Fuego under 100 mM NaCl.

Stomatal complex Cl showed a contrasting distribution pattern (Fig. 3b). Maize maintained low Cl levels under most treatments, except ES-Metronom under 50 mM NaCl. Faba bean showed higher Cl accumulation under control and CaCl₂ conditions, with Scoop reaching 0.309 atom% under control and 0.232 atom% under 50 mM CaCl₂.

Stomatal complex K differed strongly between species (Fig. 3c). Maize maintained near-zero K percentages across treatments, whereas faba bean showed higher K under control conditions and Na₂SO₄ treatment but reduced K under NaCl and CaCl₂ treatments. Fuego decreased from 0.439 atom% under control to near-zero under 100 mM NaCl.

### 3.4 Ion enrichment indices in maize and faba bean under salt stress

To compare ion distribution between stomatal complexes and bulk leaf tissue, enrichment indices for Na and Cl were calculated (Fig. 4). Positive values indicate enrichment within stomatal complexes, whereas negative values indicate depletion.

**Figure 4.**
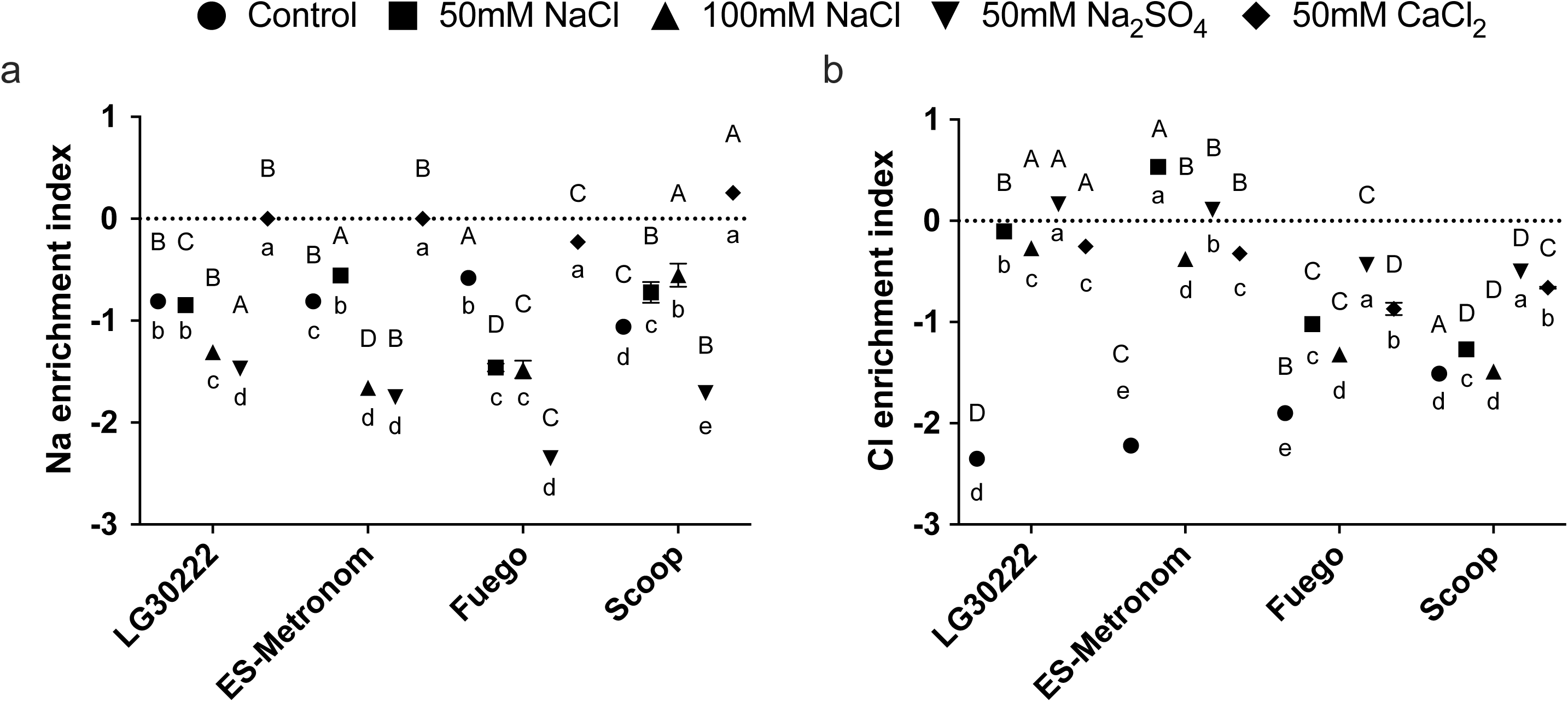
Ion enrichment indices in maize and faba bean under salt stress. **a,b**, Sodium (Na) enrichment index (a) and chloride (Cl) enrichment index (b) in two maize genotypes (LG30222 and ES-Metronom) and two faba bean genotypes (Fuego and Scoop). The enrichment index quantifies the relative accumulation or depletion of ions in the stomatal complex compared with the whole leaf. Data are presented as means ± s.e. (at least 10 stomatal complexes per treatment). Two-way ANOVA revealed significant effects of treatment, genotype and their interaction (all P < 0.001). Different uppercase letters indicate significant differences among genotypes within the same treatment, whereas lowercase letters indicate significant differences among treatments within the same genotype (Fisher’s LSD test, P < 0.05).

Na enrichment was generally negative under NaCl and Na₂SO₄ treatments, indicating lower Na representation in stomatal complexes compared with bulk leaf tissue (Fig. 4a). Maize showed strong depletion under severe Na treatments, with values reaching −1.47 and −1.75 under 50 mM Na₂SO₄ in LG30222 and ES-Metronom, respectively. Faba bean also showed Na depletion, although Scoop showed less negative enrichment than Fuego under 100 mM NaCl.

The Cl enrichment showed stronger species-dependent variation (Fig. 4b). ES-Metronom showed Cl enrichment under 50 mM NaCl (0.532), whereas LG30222 showed slight depletion under the same treatment. In contrast, faba bean showed negative Cl enrichment under NaCl treatments, with values below −1 in both genotypes. Control and CaCl₂ treatments showed negative Cl enrichment across all genotypes, but CaCl₂ resulted in significantly higher enrichment values than the control, indicating greater relative partitioning of Cl to the stomatal complex.

### 3.5 Associations between ion partitioning and gas exchange

Linear regression analyses were performed between net photosynthetic rate (A) and stomatal complex Na and Cl enrichment indices across genotypes (Fig. 5).

**Figure 5.**
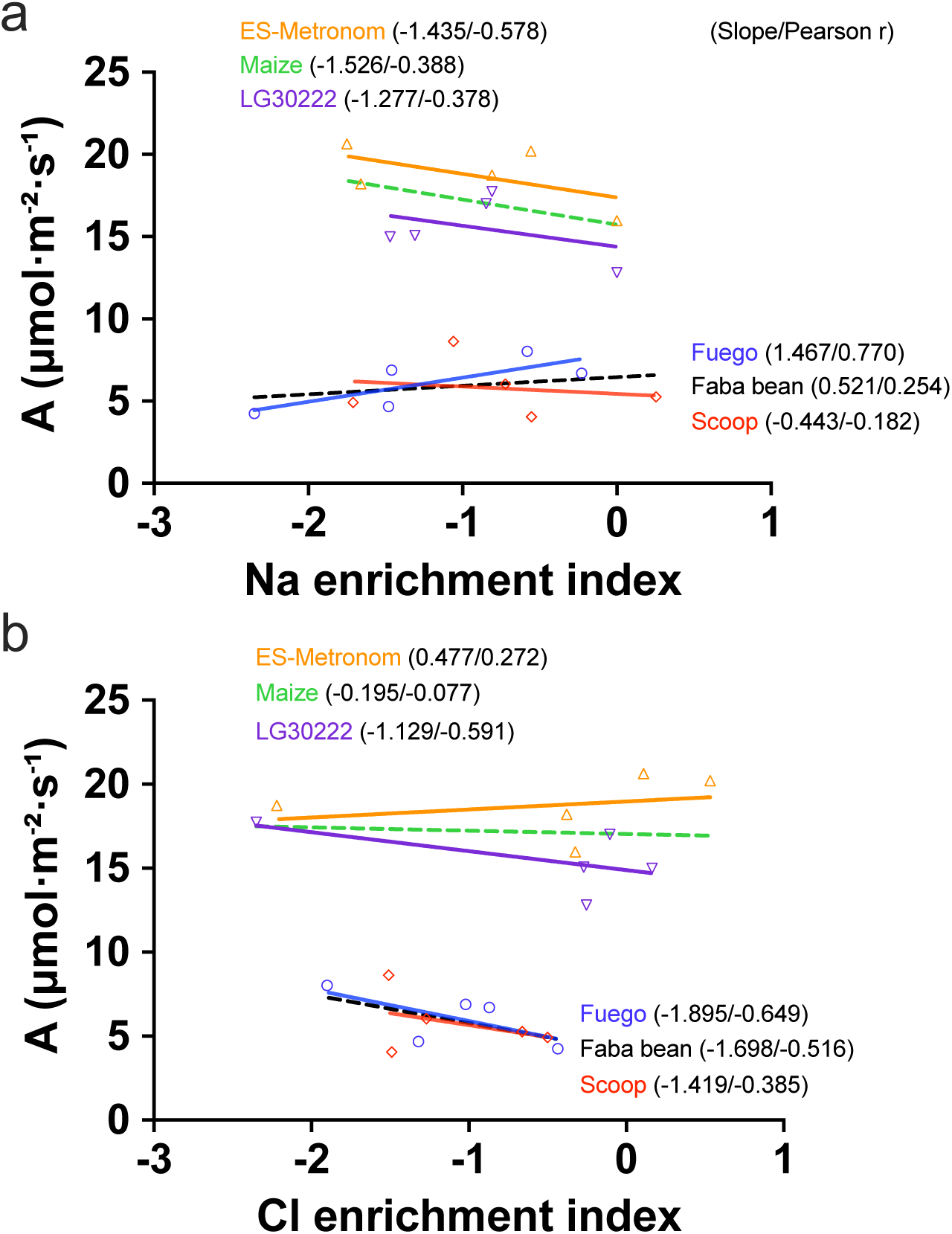
Associations between net photosynthetic rate and ion enrichment indices. **a,b**, Linear regressions between net photosynthetic rate (A) and Na (a) and Cl (b) enrichment indices across maize and faba bean genotypes. Colored lines represent genotype-specific regressions, whereas the black dashed line represents the overall regression. Numbers in parentheses denote regression slopes and Pearson correlation coefficients (r). The enrichment index quantifies the relative accumulation or depletion of ions in the stomatal complex compared with the whole leaf.

The Na enrichment showed genotype-dependent relationships with A (Fig. 5a). In maize, both genotypes showed negative correlations, with ES-Metronom showing a stronger relationship than LG30222 (r = −0.578 and −0.378, respectively). In faba bean, Fuego showed a positive correlation (r = 0.770), whereas Scoop showed a weak negative correlation (r = −0.182).

The Cl enrichment showed contrasting relationships between species (Fig. 5b). Maize showed weak and inconsistent correlations, with an overall correlation close to zero (r = −0.077). In contrast, both faba bean genotypes showed negative correlations between Cl enrichment and A, with an overall correlation of r = −0.516.

### 3.6 Multivariate integration of physiological and ionomic traits

To integrate physiological and ionomic traits, principal component analysis was performed using gas exchange parameters, leaf ion concentrations, stomatal complex ion composition and ion enrichment indices (Fig. 6). The first two principal components explained 58.8% of the total variation, with PC1 and PC2 accounting for 39.1% and 19.7%, respectively.

**Figure 6.**
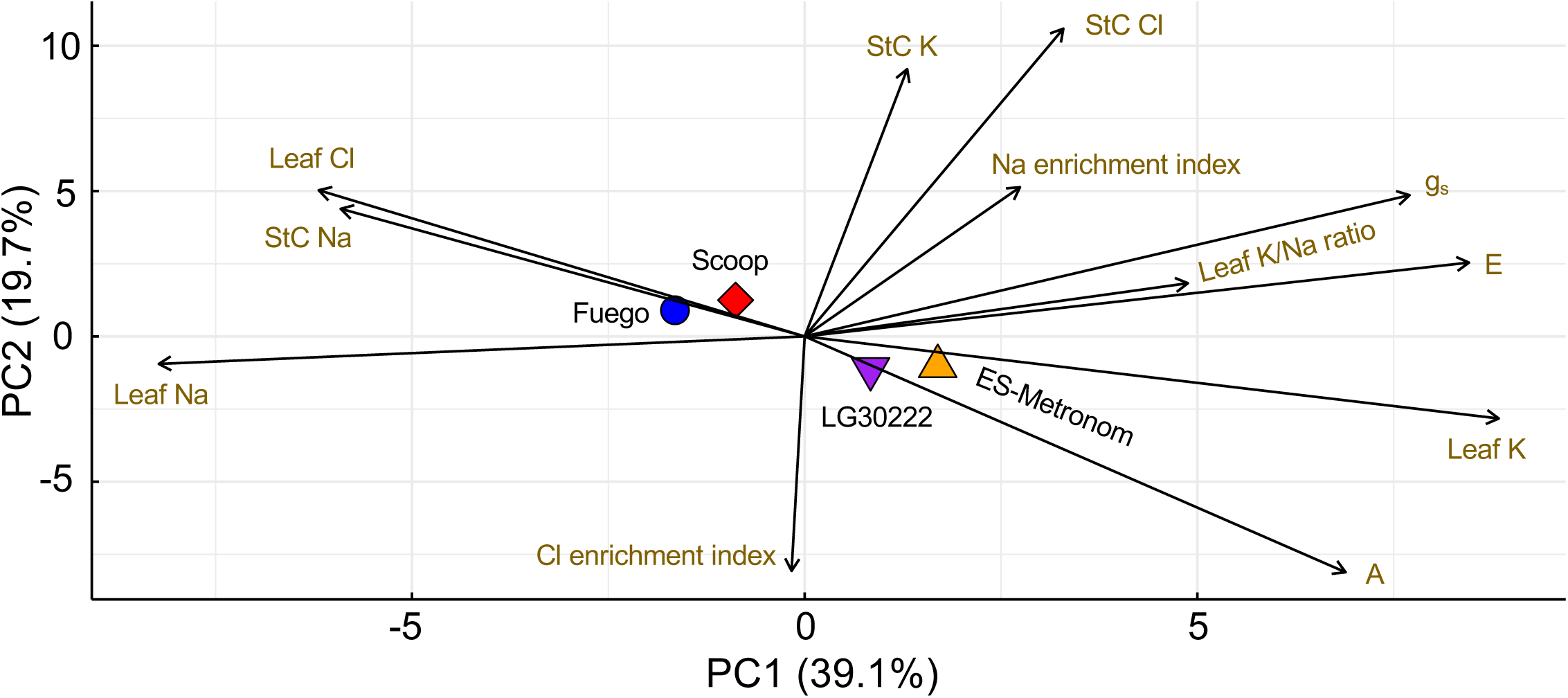
Multivariate integration of gas exchange and ionomic traits across maize and faba bean genotypes. PCA biplot showing the contribution of 12 physiological and ionomic traits to genotype separation (maize: LG30222 and ES-Metronom; faba bean: Fuego and Scoop). Traits included gas exchange parameters, leaf and stomatal complex ion traits, and ion enrichment indices, which quantify the relative accumulation or depletion of ions in the stomatal complex compared with the whole leaf. Vectors represent variable contributions and directions, whereas points indicate genotype positions within the integrated trait space.

The PCA separated maize and faba bean genotypes mainly along PC1. The maize genotypes, LG30222 and ES-Metronom, were positioned on the positive side of PC1 and showed stronger associations with gas exchange parameters (A, E and gs), leaf K and leaf K/Na ratio. In contrast, the faba bean genotypes, Fuego and Scoop, were located toward the negative side of PC1 and were associated with leaf Na, leaf Cl and stomatal complex Na.

The PC2 axis further reflected variation in ion partitioning traits. Stomatal complex K, stomatal complex Cl and Na enrichment index contributed positively to PC2, whereas Cl enrichment index showed an opposite contribution along the negative PC2 direction. The positioning of genotypes therefore reflected separation between bulk leaf ion accumulation, stomatal complex ion composition and ion enrichment patterns rather than a simple species clustering alone.

## 4 Discussion

Salinity tolerance has traditionally been interpreted through bulk leaf ion homeostasis, with reduced Na accumulation and maintenance of K considered key determinants of physiological performance (Basu et al., 2021; Garcia-Daga et al., 2025). Our results extend this framework by suggesting that gas exchange responses are associated not only with bulk leaf ion composition but also with coordinated ion partitioning between the bulk leaf and the stomatal complex. Across maize and faba bean, species-specific patterns of ion partitioning consistently accompanied contrasting physiological responses to salinity. Together, these findings identify coordinated ion partitioning as an additional organizational layer linking ion homeostasis with gas exchange regulation under salinity.

### 4.1 Species-specific ion partitioning underlies contrasting gas exchange responses to salinity

The contrasting gas exchange responses of maize and faba bean were associated with distinct ion partitioning strategies between bulk leaves and the stomatal complex. Maize maintained higher photosynthetic rates and stomatal conductance under salinity, accompanied by lower leaf Na accumulation, greater K retention and higher K/Na ratios than faba bean (Figs. 1 and 2). These differences are consistent with previous studies showing that maize restricts Na and Cl accumulation in photosynthetic tissues and maintains ionic homeostasis more effectively under salinity (Abd El-Samad et al., 2011; Iqbal et al., 2020), whereas faba bean exhibits greater Na accumulation in both apoplastic and symplastic compartments, contributing to ionic imbalance and reduced photosynthetic performance (Shahzad et al., 2013; Tavakkoli et al., 2010). Extending these previous observations, our results demonstrate that species-specific ion regulation also occurs at the stomatal complex level, where maize maintained lower Na abundance and stronger Na depletion relative to bulk leaf tissue than faba bean under saline conditions (Figs. 3 and 4). Thus, salt tolerance is associated not only with limiting total ion accumulation but also with precise spatial regulation of ions within functional leaf compartments especially in photosynthetically relevant organs like stomata.

This finding extends the classical view that salinity tolerance depends primarily on Na exclusion and K retention (de Souza Mateus et al., 2019; Niu et al., 2018) by highlighting spatial ion allocation as an additional regulatory layer. In maize, increased Na enrichment in the stomatal complex was associated with reduced photosynthetic performance, whereas faba bean showed no comparable negative relationship (Fig. 5a). This suggests that stomatal complex Na partitioning is more closely linked with photosynthetic regulation in maize, whereas photosynthetic limitation in faba bean likely involves multiple ion-related constraints. Consistently, the PCA associated maize with gas exchange, leaf K and K/Na ratio, whereas faba bean was associated with leaf and stomatal complex Na and Cl accumulation in opposition to gas exchange performance (Fig. 6). Together, these results demonstrate that species-specific ion partitioning contributes to divergent physiological responses to salinity.

### 4.2 Stomatal complex ion partitioning provides mechanistic insight beyond bulk leaf ionomics

Bulk leaf ion concentrations are widely used to evaluate salinity tolerance; however, they provide limited information about the spatial distribution of elements within functional tissues like guard cells and stomata. Previous studies have demonstrated that salinity tolerance involves multiple components beyond bulk Na accumulation, including tissue tolerance and intracellular ion compartmentation (Munns et al., 2016; Rajendran et al., 2009). Although maize and faba bean differed strongly in leaf Na, Cl and K accumulation (Fig. 2), our enrichment analysis revealed that the relative allocation of elements between bulk leaf tissue and the stomatal complex provided additional information associated with gas exchange performance (Fig. 4). Under NaCl and Na₂SO₄ stress, Na was generally depleted from the stomatal complex relative to bulk leaf tissue, with stronger depletion observed in maize than in faba bean under severe Na stress (Fig. 4a). This pattern was consistent with the maintenance of higher photosynthetic activity and stomatal conductance in maize, whereas altered Na partitioning in faba bean coincided with greater physiological impairment (Figs. 1 and 5a).

The enrichment index therefore provides a quantitative perspective on elemental distribution beyond bulk leaf elemental concentrations. This distinction is important because total leaf accumulation does not reveal how elements are partitioned among cell types with different physiological sensitivities. Cell-specific elemental compartmentation has emerged as an important mechanism of salinity adaptation. For example, salt tolerant halophytes sequester Na and Cl into epidermal bladder cells, reducing ion accumulation in photosynthetically active mesophyll tissues and maintaining cellular homeostasis (Agarie et al., 2007; Kiani-Pouya et al., 2017). Similarly, differential Na distribution between epidermal and mesophyll cells contributes to contrasting salt tolerance in cotton by limiting Na exposure in sensitive cellular compartments (Peng et al., 2016). Extending these concepts, our results identify the stomatal complex as an additional functional compartment where elemental partitioning is associated with gas exchange regulation. The PCA further supports this interpretation, showing that stomatal complex elemental traits and enrichment indices captured additional variation beyond bulk leaf elemental traits (Fig. 6).

### 4.3 Na and Cl exhibit distinct partitioning patterns associated with species-specific salinity responses

Although Na and Cl are often considered together as major components of salinity stress, our results indicate that these elements exhibit distinct partitioning patterns and may contribute differently to physiological responses. Maize salt tolerance is commonly associated with efficient Na exclusion from shoots and/or sequestration into vacuoles, which limits Na accumulation in photosynthetically active tissues (Rizk et al., 2024; Zhang et al., 2018). In contrast, faba bean is more prone to Na accumulation in leaf tissues and apoplastic compartments, where excessive Na disrupts ionic homeostasis and impairs physiological function (Percey et al., 2014; Shahzad et al., 2013). Extending these previous observations, our results reveal that species-specific Na regulation also occurs within the stomatal complex. Maize maintained lower stomatal complex Na abundance and stronger Na depletion relative to bulk leaf tissue under severe salt stress, whereas faba bean showed higher stomatal complex Na representation and weaker depletion (Figs. 3 and 4a). These contrasting partitioning patterns were associated with different relationships between Na enrichment and photosynthesis, indicating that the physiological impact of Na depends not only on total accumulation but also on its spatial distribution within functional leaf compartments (Fig. 5a).

In contrast, Cl displayed a distinct partitioning behavior between species. Maize generally limits Cl accumulation in photosynthetically active tissues through restricted root-to-shoot transport and sequestration in roots and leaf sheaths, thereby maintaining physiological function under chloride stress (Zhang et al., 2020; Zhang et al., 2019). In faba bean, excessive Cl translocation and accumulation in leaves have been associated with reduced photosynthetic capacity and increased salt sensitivity (Franzisky et al., 2019; Richter et al., 2019). Consistent with these contrasting strategies, maize showed relatively limited changes in Cl enrichment across treatments, whereas faba bean exhibited stronger Cl depletion from the stomatal complex under NaCl stress (Fig. 4b). Moreover, Cl enrichment was negatively associated with photosynthetic performance in faba bean, suggesting that altered Cl partitioning contributed to reduced physiological performance under salt stress (Fig. 5b). Together with the PCA analysis, which showed distinct contributions of Na- and Cl-related traits to multivariate variation (Fig. 6), these results indicate that Na and Cl represent functionally distinct components of salinity stress. Their effects on physiological performance are determined not only by their total accumulation but also by their compartment-specific allocation and interaction with elemental homeostasis processes.

## 5 Conclusion

Our study demonstrates that salinity tolerance is associated not only with the capacity to restrict ion accumulation at the whole leaf level but also with the spatial partitioning of ions between the bulk leaf and the stomatal complex. The contrasting gas exchange responses of maize and faba bean were closely linked to species-specific patterns of Na, Cl and K distribution, highlighting the importance of compartment-specific ion regulation in maintaining physiological function under salinity. By integrating bulk leaf ionomics with stomatal complex ion profiling, we identify the stomatal complex as a functionally distinct ionomic compartment that provides additional insight beyond conventional leaf ion measurements. These findings establish stomatal complex ion partitioning as an important component of salinity adaptation and provide a framework for understanding the cellular basis of species-specific salt tolerance.

## Acknowledgement

The work was financially supported by the Deutsche Forschungsgemeinschaft (DFG, German Research Foundation, ZO 118/15–1), project number 491678431. We thank Dr. Claus J. Burkhardt and Dr. Birgit Schröppel at NMI Natural and Medical Sciences Institute, University of Tübingen for technical support with cryo-SEM-EDX system and data analysis. We also thank Ms. Christiane Beierle and Ms. Dagmar Repper for cryo-specimen preparation.

## Author Contributions

X.D.Z. led the conceptualization of the study, developed the methodology, performed the investigation, curated and analyzed the data, developed software and visualizations, administered the project, and wrote the original draft of the manuscript as well as the revised versions. G.H.W. contributed to the investigation by supporting experimental work. C.Z. contributed to the conceptualization of the study, secured funding, administered the project, and reviewed and edited the manuscript. All authors read and approved the final manuscript and agree to be accountable for all aspects of the work, ensuring that questions related to the accuracy or integrity of any part of the research are appropriately addressed.

## AI use

During the preparation of this manuscript, AI-assisted technology was used solely to improve language clarity and readability. After using this tool/service, the authors reviewed and edited the content as needed and take full responsibility for the content of the published article.

