## Supplementary files for "Stomatal Complex Ionomes Explain Divergent Gas Exchange Responses to Salinity in Maize and Faba Bean"

### Supplementary Method S1. Cryo-SEM-EDX analysis of stomatal complex surface elemental composition

Cryo-SEM-EDX was used to characterize the surface-resolved distribution of mobile elements, particularly Na, K and Cl, across stomatal complex cells of maize leaves. To preserve native elemental distributions, conventional chemical fixation and embedding procedures were avoided because they may introduce external ions or alter soluble elemental pools. Epidermal tissues were plunge-frozen in liquid nitrogen to maintain the native ion distribution state (Franzisky et al., 2025).

Epidermal strips from the same fifth fully expanded leaves used for gas exchange measurements were mounted on copper grids and transferred into a cryo-preparation chamber (Quorum Technologies) connected to a Zeiss Crossbeam 550 SEM. No freeze fracture or sublimation treatment was applied to avoid redistribution or loss of mobile ions. Samples were sputter-coated with platinum (5 mA, 45 s; approximately 35 nm) to reduce charging effects and improve X-ray signal stability. SEM imaging and EDX measurements were performed at an accelerating voltage of 8 kV, beam current of 1 nA, magnification of 2126× and working distance of 5.5 mm. The sample stage was tilted by 45° to improve X-ray collection efficiency. Because stomatal complexes possess complex three-dimensional surface structures, cells with optimal orientation relative to the detector were manually selected to minimise shadowing effects and improve measurement consistency.

Elemental spectra were acquired using an Oxford Instruments X-MaxN 150 SDD detector controlled by AZtec software (version 5.1). Spectral fitting included biological elements (C, N, O, Na, Mg, P, S, Cl, K and Ca) together with instrumental signals derived from the copper support grid, platinum coating and gold sample holder. Quantification was performed using the AZtec standardless routine, including background subtraction, peak deconvolution, absorption correction, fluorescence correction and pulse pile-up correction. Hydrogen was not detectable by EDX. Carbon and oxygen were retained during normalization as dominant structural elements of plant tissues. Nitrogen was included during spectral fitting to improve peak separation but excluded from quantitative interpretation because the low-energy N Kα signal is strongly affected by absorption, background interference and coating effects.

At the applied accelerating voltage of 8 kV, the electron interaction volume extended to approximately 1.5 µm in depth. Therefore, EDX signals primarily represent the elemental composition of the outer surface region of epidermal cells, including the cuticle and outer cell wall-associated ion pools, rather than intracellular ion concentrations. Accordingly, atom% values obtained from cryo-SEM-EDX were interpreted as relative surface elemental composition and not as absolute cellular ion contents. This surface-resolved analysis provides complementary information to bulk leaf ion concentrations determined by ICP-OES, enabling assessment of relative ion distribution and enrichment within stomatal complexes.

Element selection for biological interpretation was based on both spectral reliability and biological relevance. Under the applied measurement conditions, Na, Mg, P, S, Cl, K and Ca showed sufficient excitation conditions for reliable detection, whereas elements requiring higher excitation energies, including Mn, Fe, Zn, Mo and Co, were excluded from quantitative analysis. Representative EDX spectra from all genotype–treatment combinations are presented in Figure S1. Based on clear characteristic peaks and consistent detection across samples, Na, K and Cl were selected for comparative analysis of salt-induced elemental distribution patterns.

Regions of interest (ROIs) corresponding to the entire stomatal complex were manually defined based on SEM morphology. In faba bean, the stomatal complex included the stomatal pore and the surrounding pair of guard cells, whereas in maize it additionally included the pair of subsidiary cells surrounding the guard cells. In this study, the stomatal complex was treated as a single functional unit, and elemental signals from individual component cells were not separated. At least 10 stomatal complexes per treatment were analysed, selected from five epidermal peel samples obtained from the same fifth leaves used for gas exchange measurements. Each spectrum was acquired for at least 5 min. Elemental abundances were expressed as atomic percentages (atom%) and normalized to the sum of reliably detected biological elements (C, O, Na, Mg, P, S, Cl, K and Ca) for semi-quantitative comparison.

$$Atom\%_{\ normalized} = \frac{Atom\%_{element}}{\sum(C + O + Na + Mg + P + S + Cl + K + Ca)} \times 100$$

Because EDX detects characteristic X-ray emissions from the near-surface interaction volume, measured values reflect relative elemental composition of the analyzed surface region rather than total elemental accumulation within the whole cell. When elemental signals, such as Na or Cl in sensitive genotypes, were indistinguishable from background noise and lacked clear characteristic peaks, they were considered below the detection limit and assigned a value of zero for statistical analysis and visualization. These criteria enabled consistent comparison of relative ion enrichment patterns among genotypes and salt treatments.

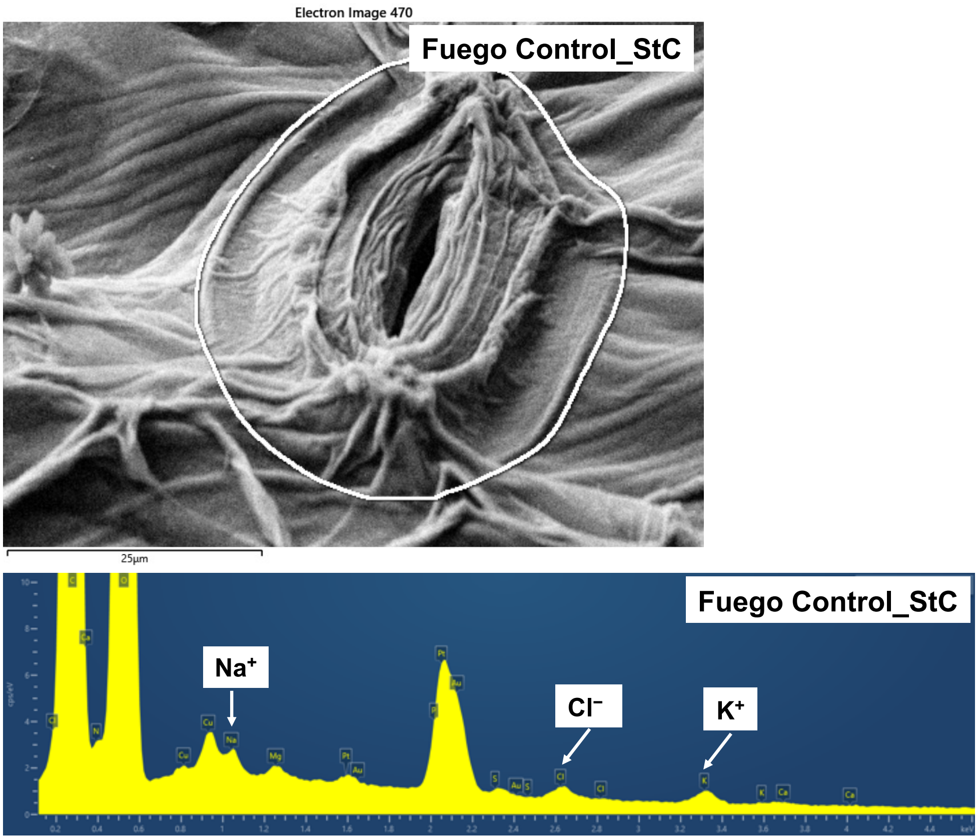

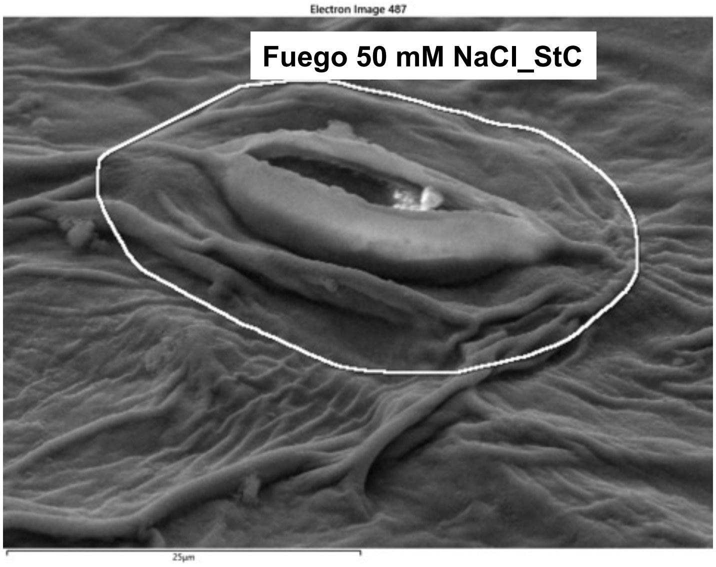
Electron Image 487
Fuego 50 mM NaCl_StC
25µm

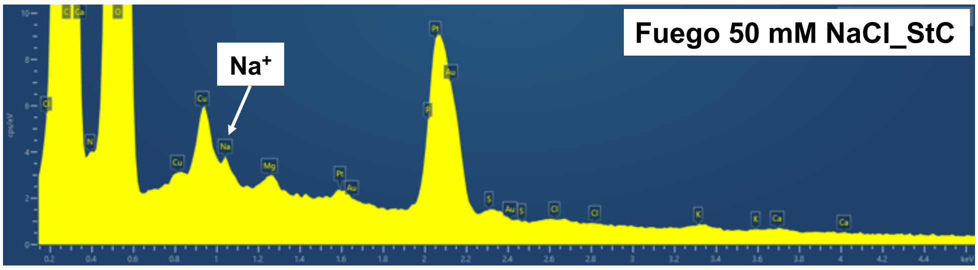
Na+
Fuego 50 mM NaCl_StC

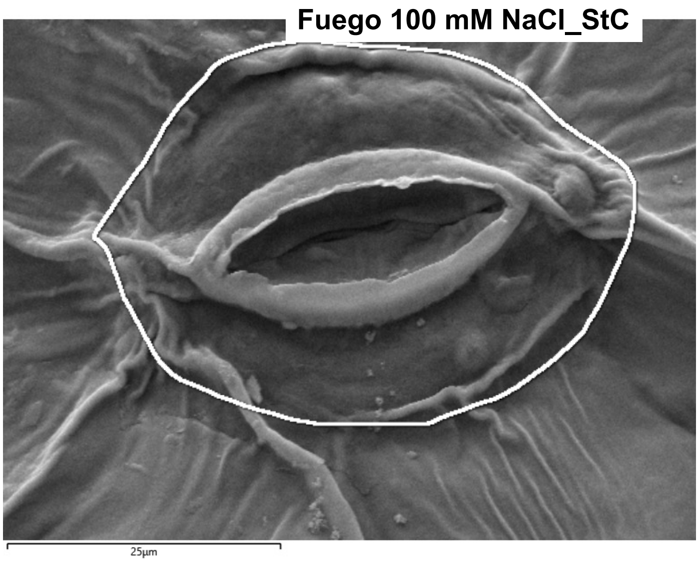
Fuego 100 mM NaCl_StC
25µm

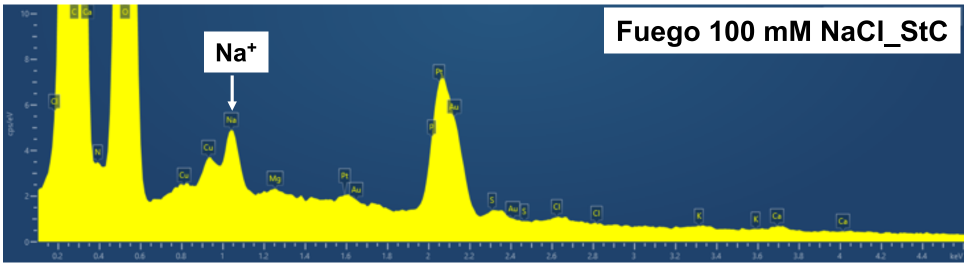
Na+
Fuego 100 mM NaCl_StC

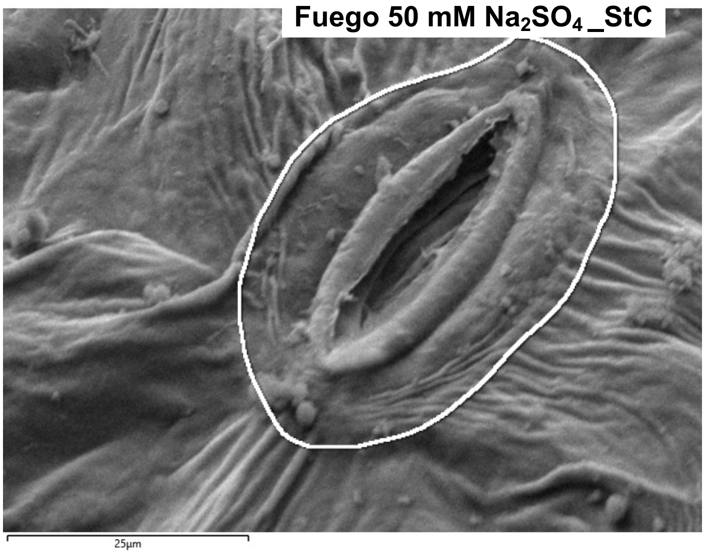
Fuego 50 mM $Na_2SO_4$_StC
25µm

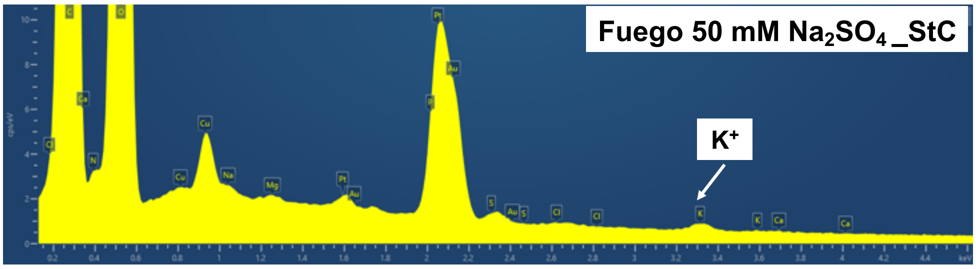
Fuego 50 mM $Na_2SO_4$_StC
$K^+$

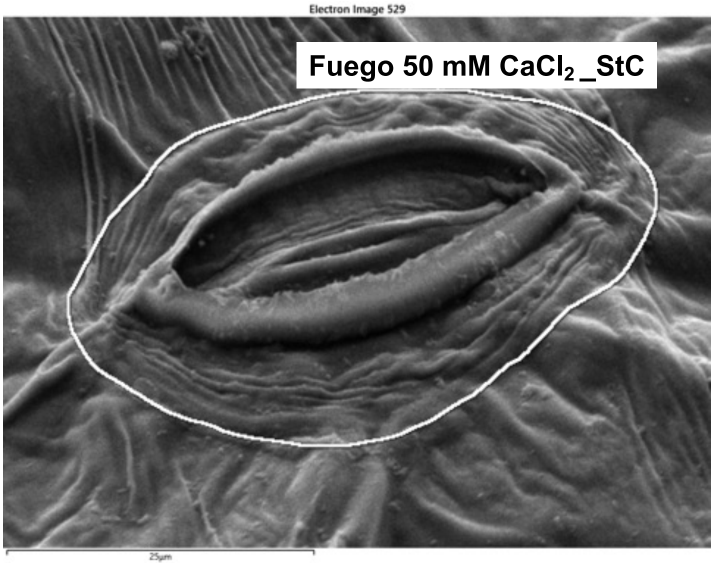
Electron Image 529
Fuego 50 mM $CaCl_2$_StC
25µm

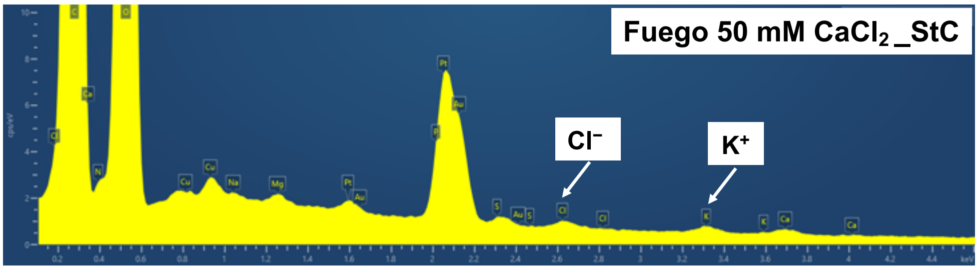
Fuego 50 mM $CaCl_2$_StC
$Cl^-$
$K^+$

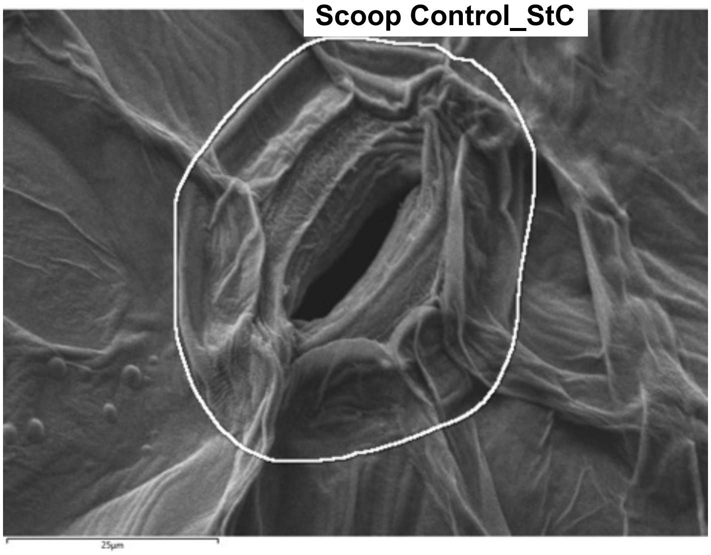
Scoop Control_StC

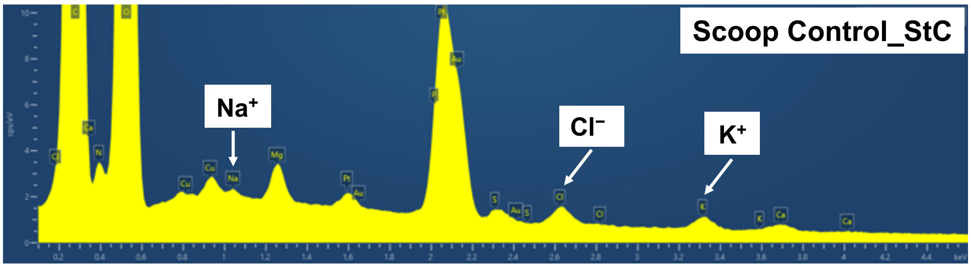
Scoop Control_StC
Na+
Cl−
K+

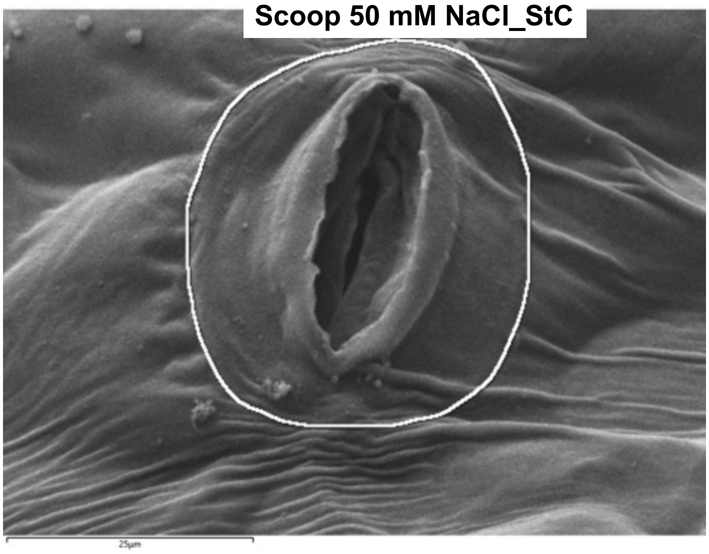
Scoop 50 mM NaCl_StC

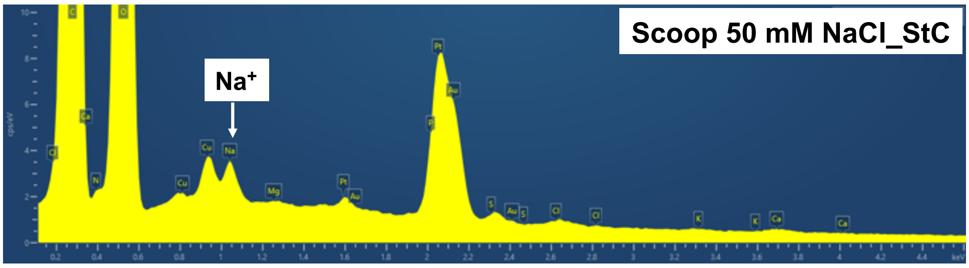
Scoop 50 mM NaCl_StC
Na+

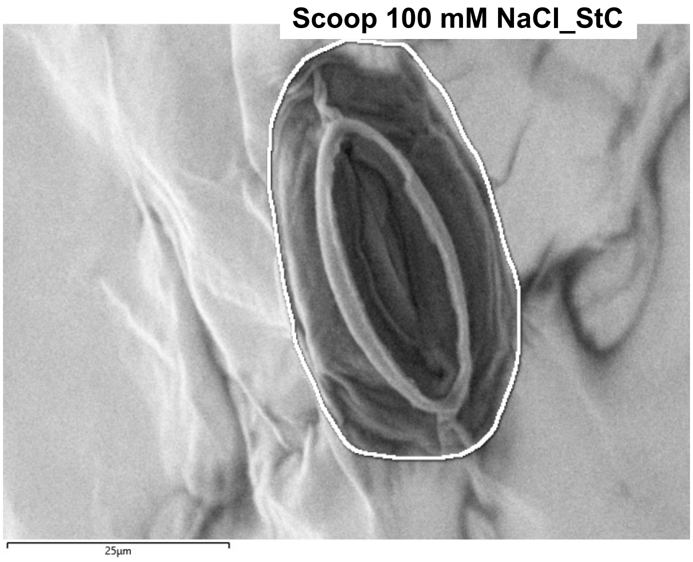
Scoop 100 mM NaCl_StC
25µm

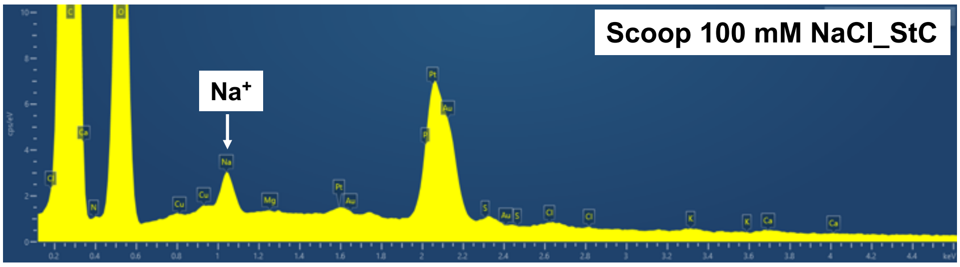
Scoop 100 mM NaCl_StC
Na+

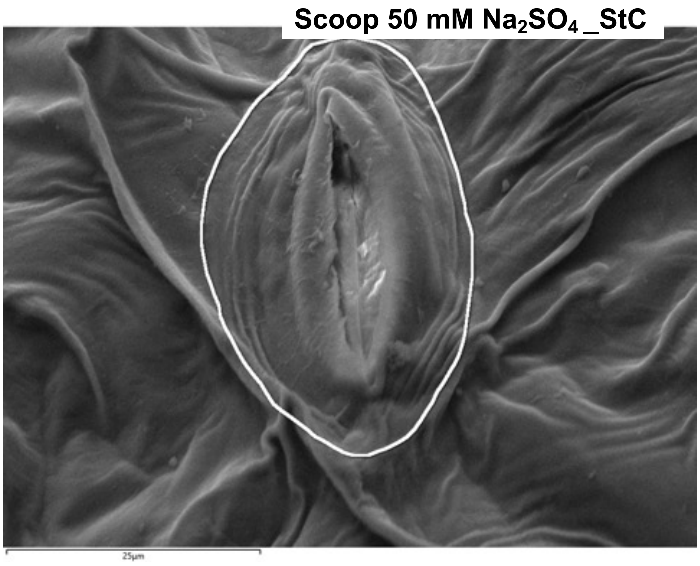
Scoop 50 mM Na2SO4 _StC
25µm

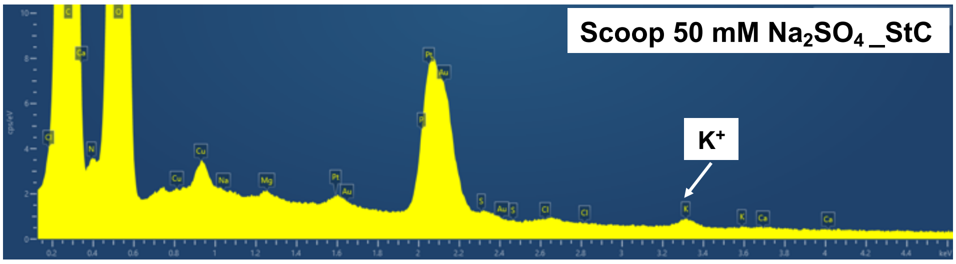
Scoop 50 mM Na2SO4 _StC
K+

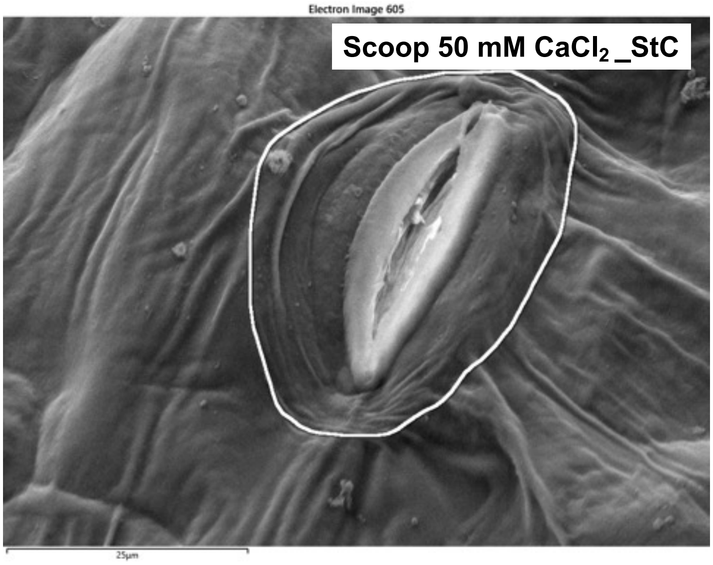
Electron Image 605
Scoop 50 mM $CaCl_2$ _StC
25µm

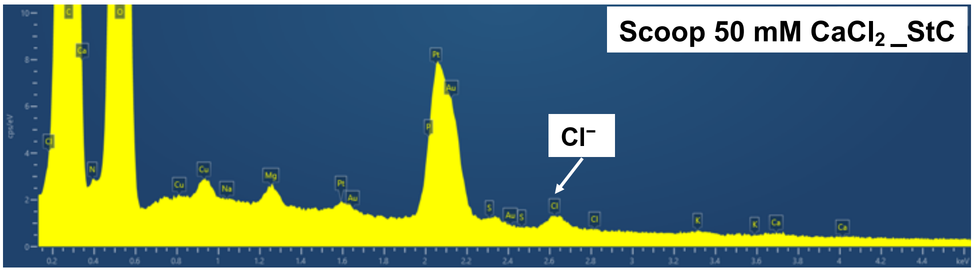
Scoop 50 mM $CaCl_2$ _StC
$Cl^-$

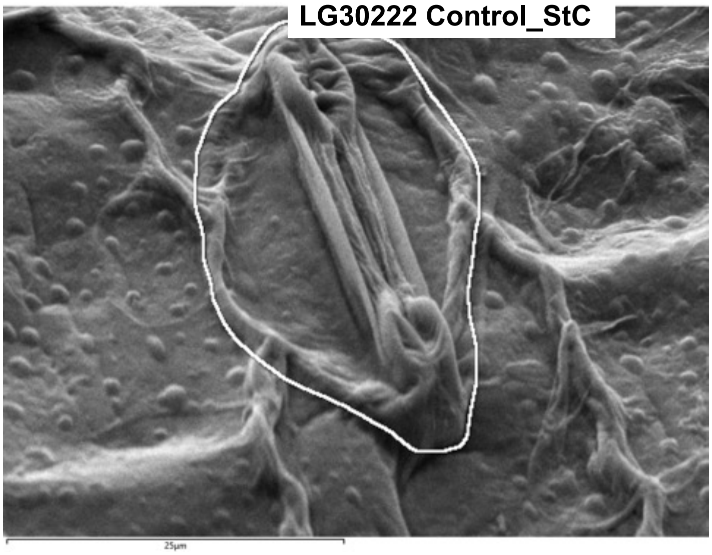
LG30222 Control_StC
25µm

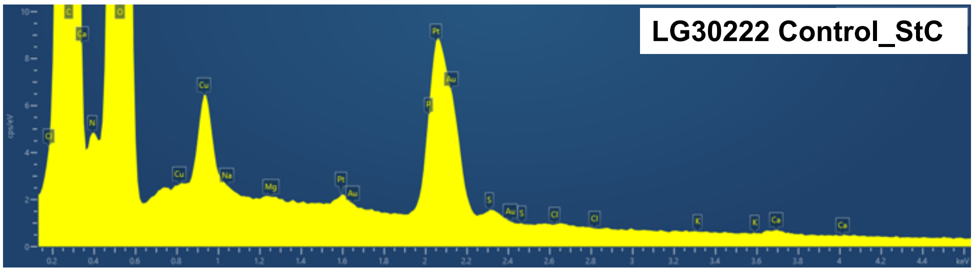
LG30222 Control_StC

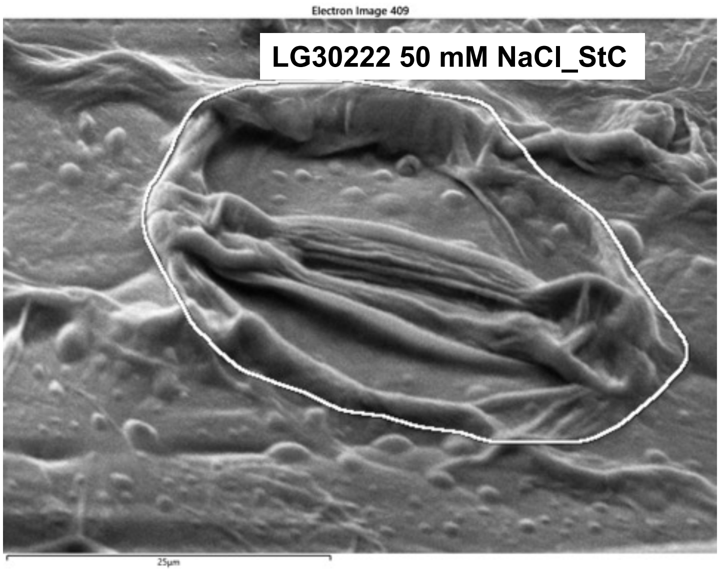
Electron Image 409
LG30222 50 mM NaCl_StC
25µm

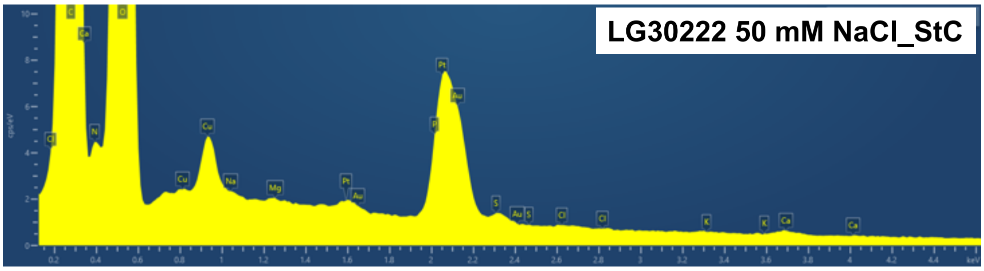
LG30222 50 mM NaCl_StC

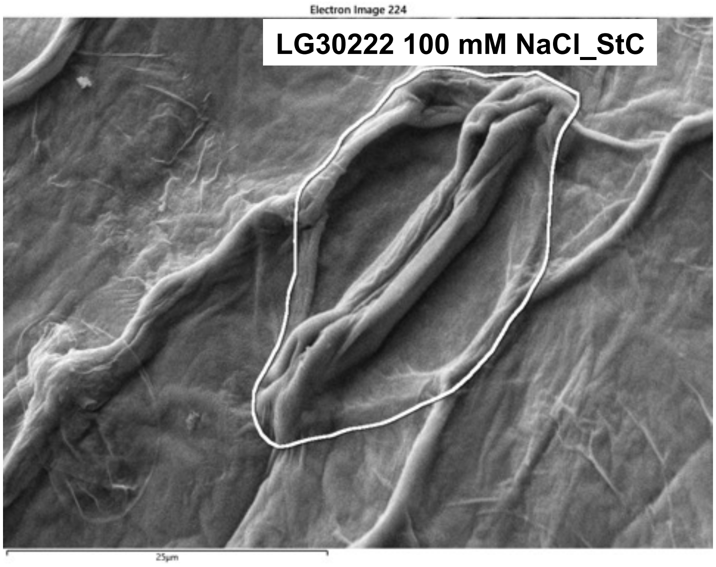
Electron Image 224
LG30222 100 mM NaCl_StC
25µm

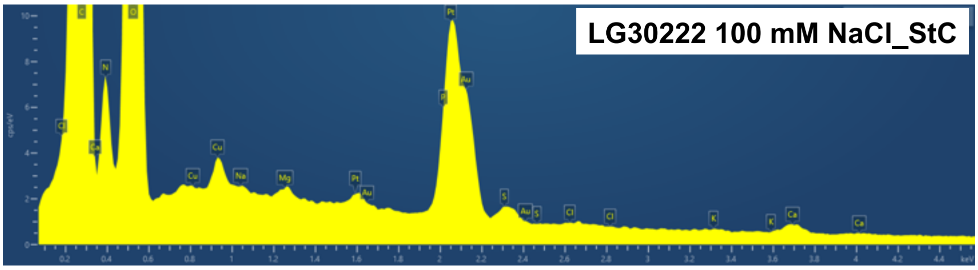
LG30222 100 mM NaCl_StC

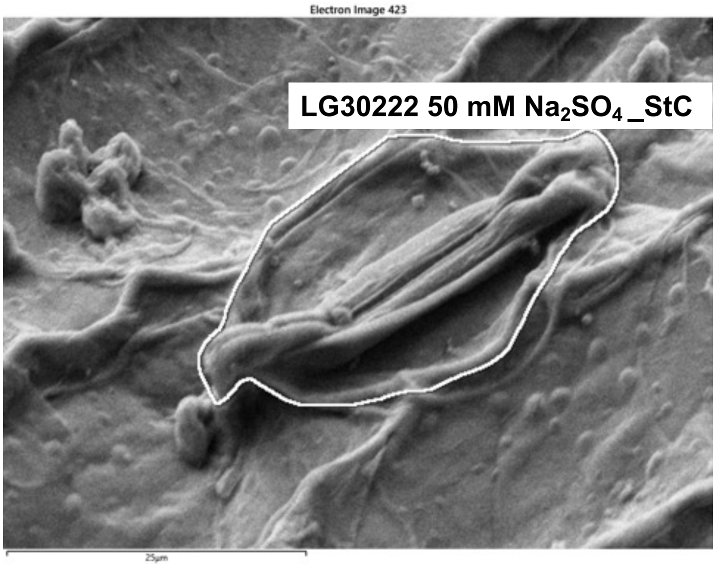
Electron Image 423
LG30222 50 mM $Na_2SO_4$_StC
25µm

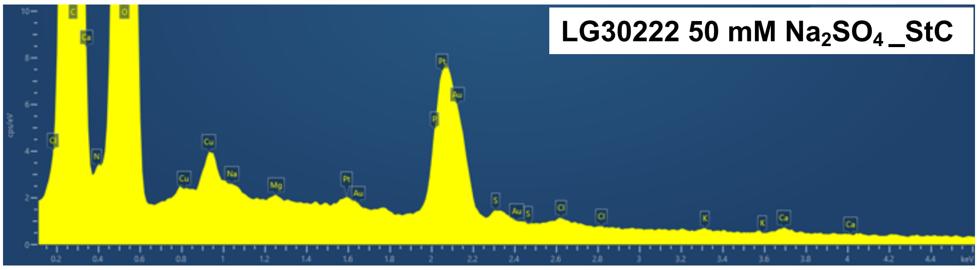
LG30222 50 mM $Na_2SO_4$_StC

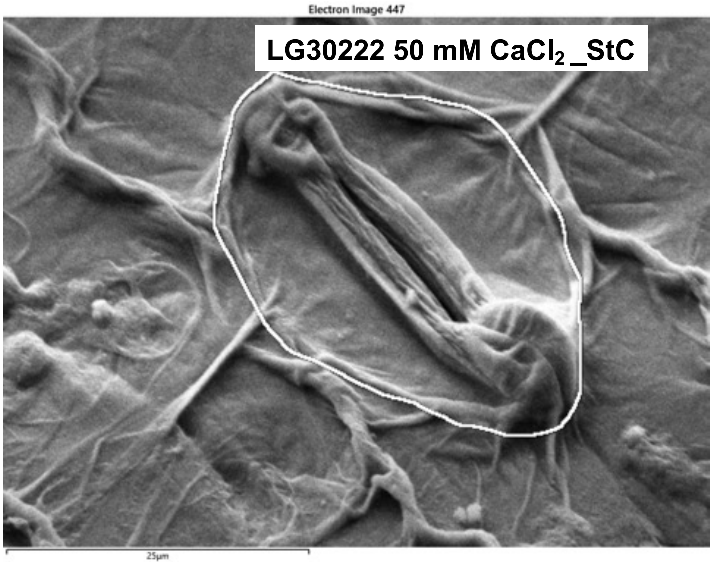
Electron Image 447
LG30222 50 mM $CaCl_2$_StC
25µm

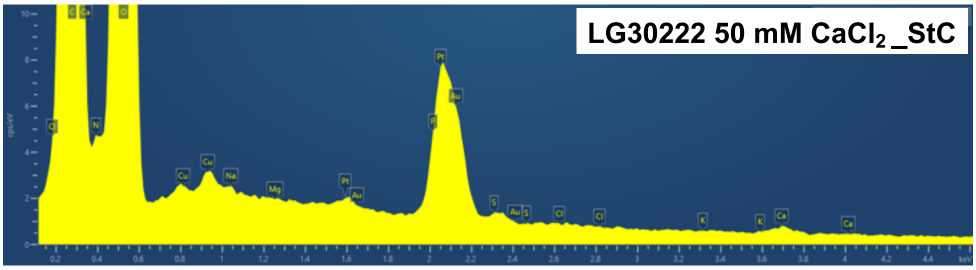
LG30222 50 mM $CaCl_2$_StC

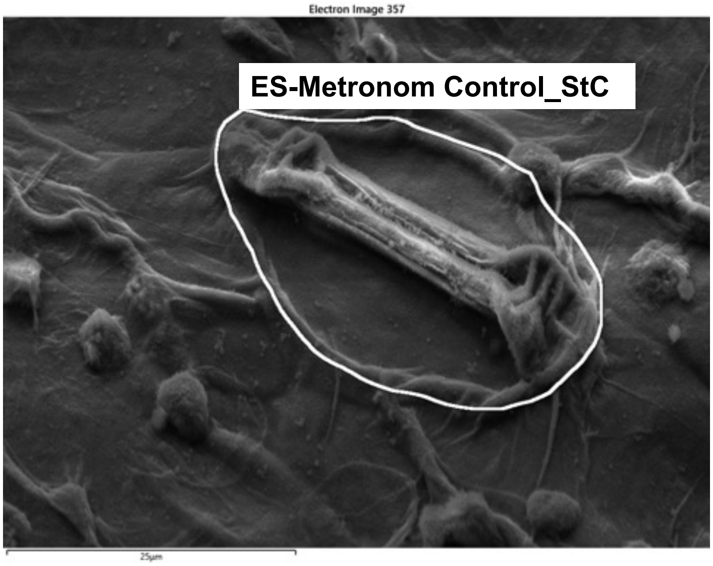
Electron Image 357
ES-Metronom Control_StC
25µm

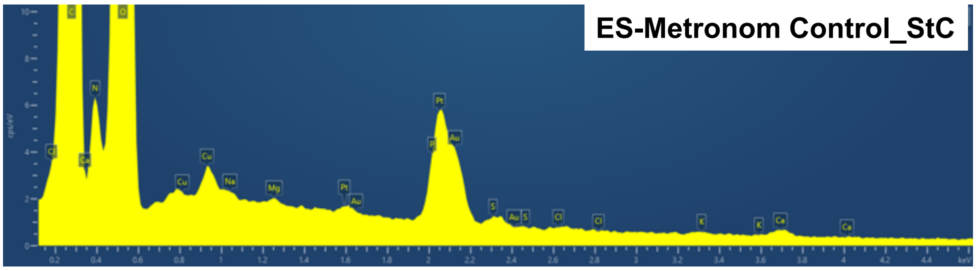
ES-Metronom Control_StC

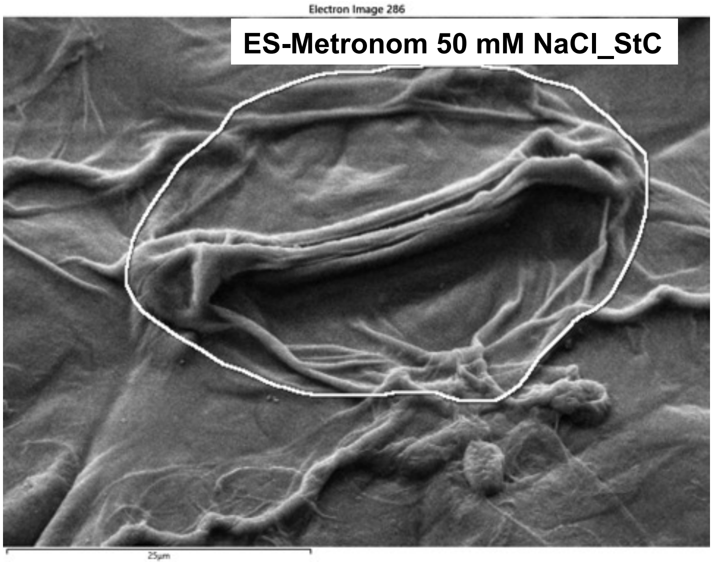
Electron Image 286
ES-Metronom 50 mM NaCl_StC
25µm

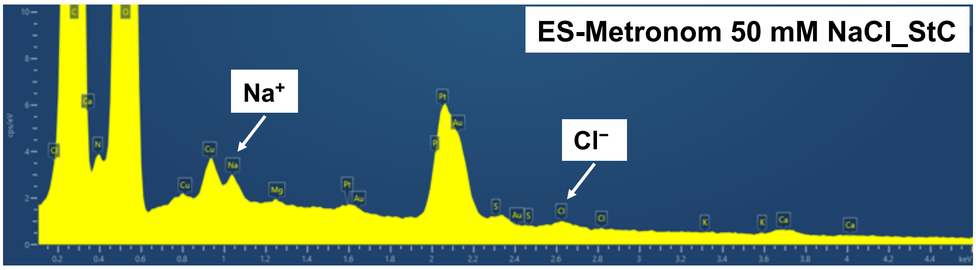
ES-Metronom 50 mM NaCl_StC
Na+
Cl−

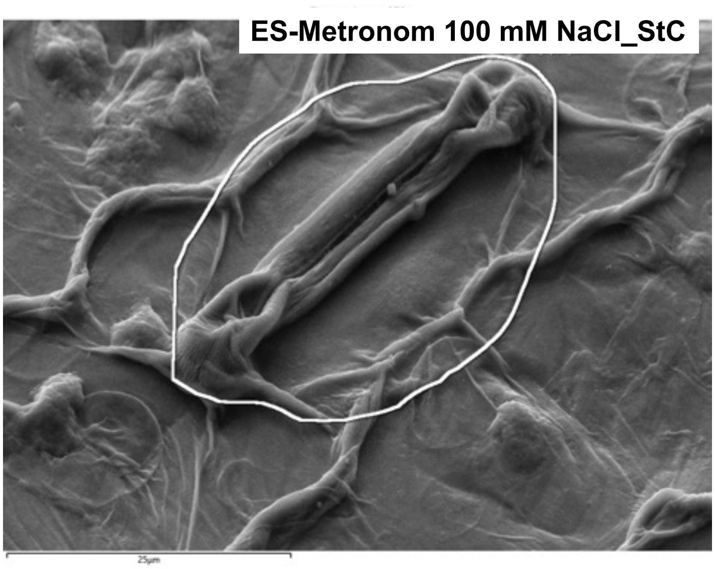
ES-Metronom 100 mM NaCl_StC

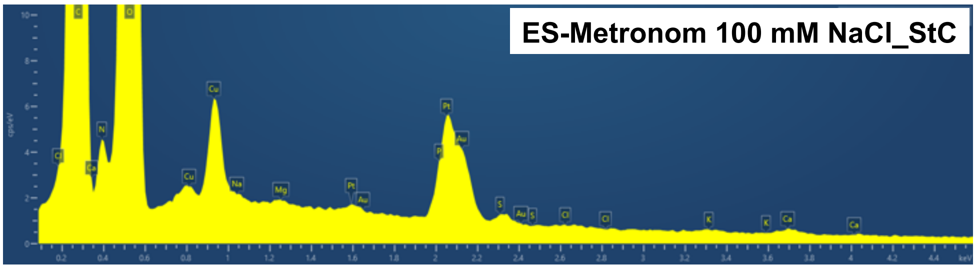
ES-Metronom 100 mM NaCl_StC

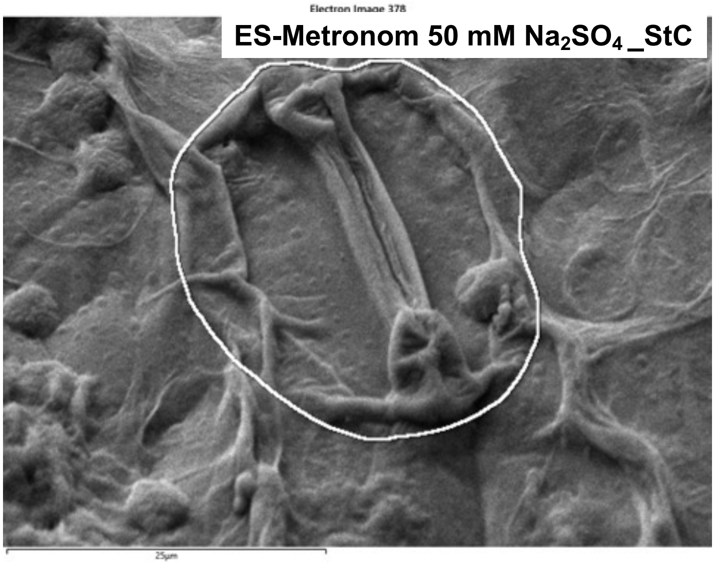
Electron Image 378
ES-Metronom 50 mM $Na_2SO_4$_StC

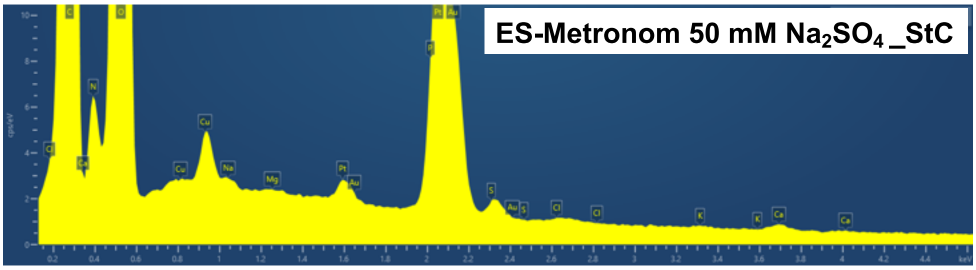
ES-Metronom 50 mM $Na_2SO_4$_StC

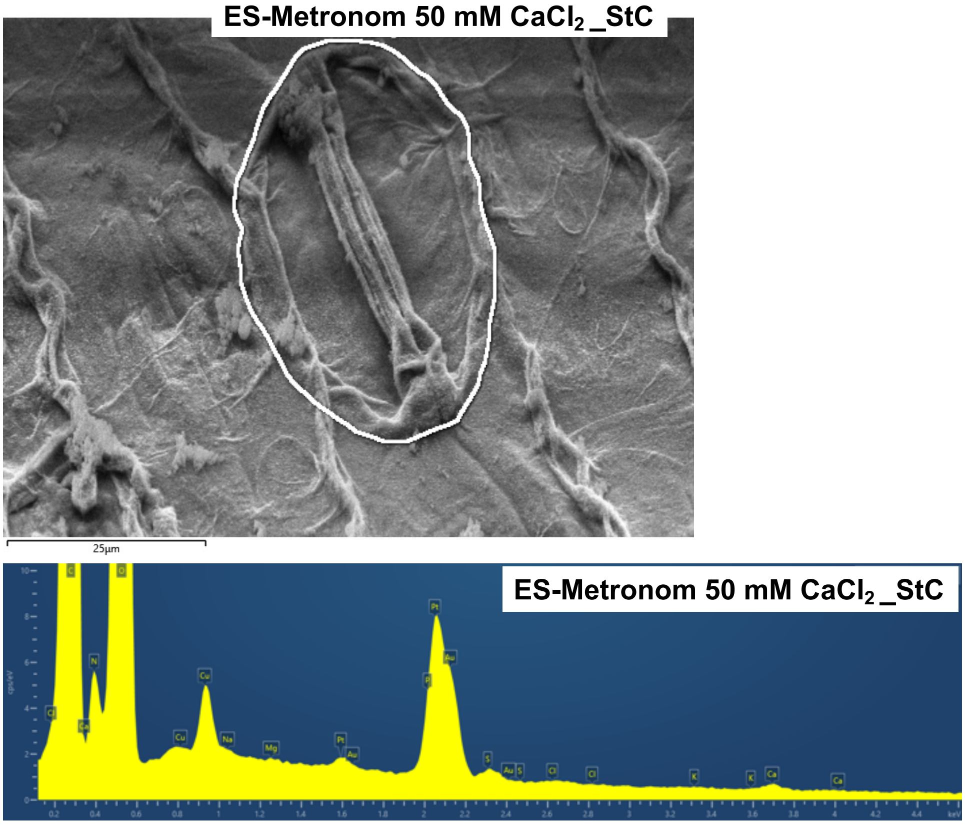


**Supplementary Figure S1. Representative cryo-SEM-EDX spectra of stomatal complex surface regions across maize and faba bean genotypes and salt treatments**

Representative energy-dispersive X-ray spectroscopy (EDX) spectra obtained from stomatal complex surface regions of two contrasting maize genotypes (LG30222 and ES-Metronom) and two contrasting faba bean genotypes (Fuego and Scoop) under control and salt treatments. Spectra from all genotype–treatment combinations are shown to demonstrate elemental detection reliability across experimental conditions. Characteristic peaks corresponding to Na, K and Cl are specifically labelled because these elements were selected for comparative analysis of salt-induced elemental distribution patterns. Other detected elements, including Mg and Ca, were retained during spectral fitting and normalization and are shown only through automatic peak annotation in the exported spectra. Signals originating from the copper support grid (Cu), platinum coating (Pt) and gold sample holder (Au) are also automatically annotated. EDX signals represent relative elemental composition of the near-surface region, with a detection depth of approximately 1.5 µm, rather than intracellular ion concentrations.

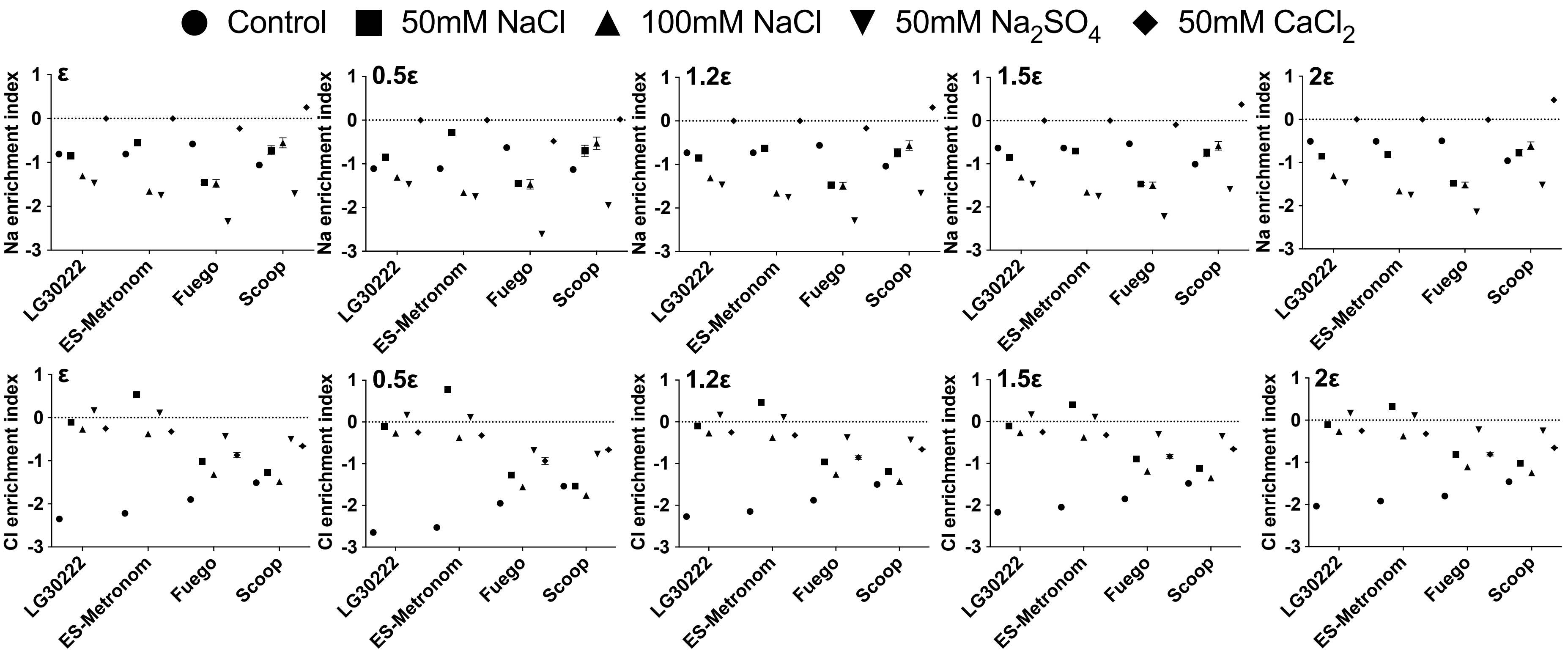


**Supplementary Figure S2. Sensitivity analysis of pseudocount selection for Na and Cl enrichment indices across maize and faba bean genotypes**

Sensitivity analysis showing the effect of different pseudocount values (ε) on Na and Cl enrichment indices calculated from stomatal complex surface elemental signals relative to corresponding whole-leaf ion concentrations across two maize genotypes (LG30222 and ES-Metronom) and two faba bean genotypes (Fuego and Scoop). The reference ε value was defined as half of the minimum non-zero value across all samples, and alternative values corresponding to 0.5×, 1×, 1.2×, 1.5× and 2× of this value were evaluated. Similar treatment- and genotype-associated enrichment patterns were observed across the tested ε values, supporting the selection of the reference pseudocount for enrichment index calculation. The enrichment index was calculated as the $\log_{10}$ ratio between normalized stomatal complex surface elemental signals and corresponding normalized whole-leaf ion concentrations.
